# Coincident Fluorescence Burst Analysis of dUTP-Loaded Exosome-Mimetic Nanovesicles

**DOI:** 10.1101/2021.10.11.463914

**Authors:** Maryam Sanaee, Elin Sandberg, K. Göran Ronquist, Jane M. Morrell, Jerker Widengren, Katia Gallo

## Abstract

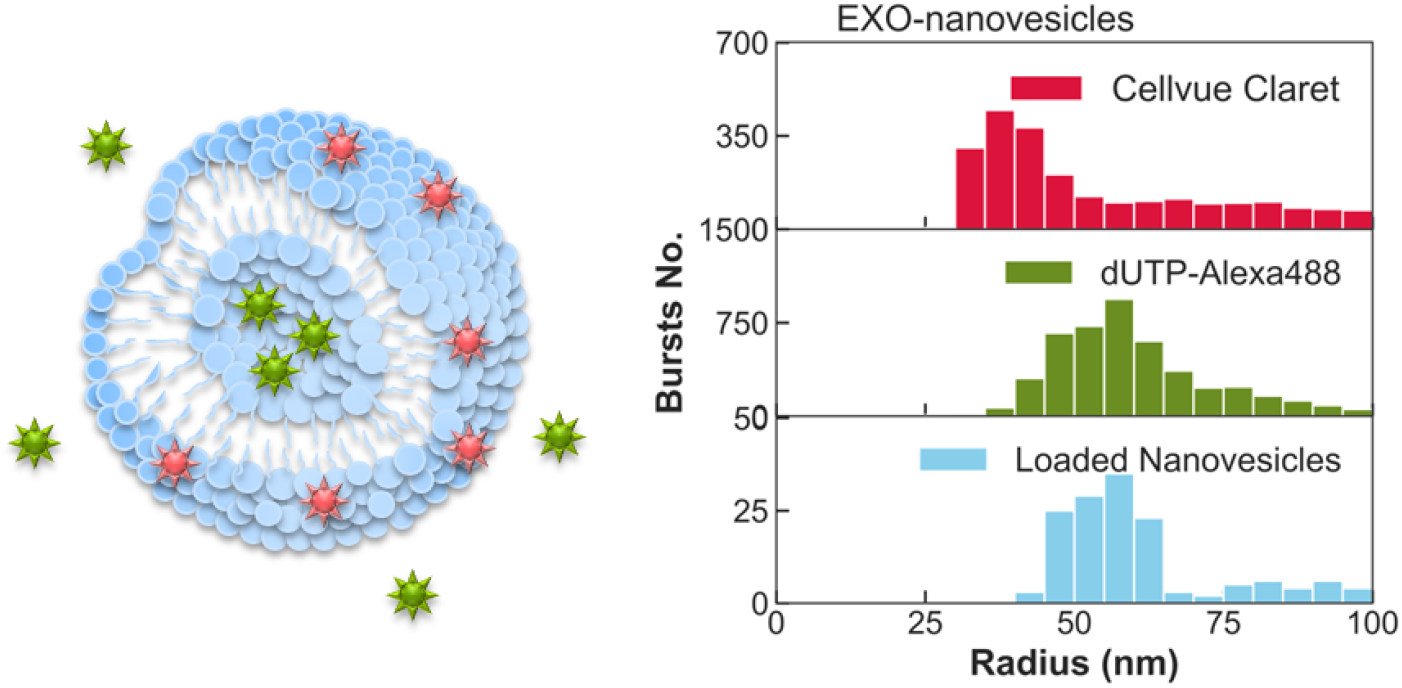

The targeting functionality and low immunogenicity of exosomes and exosome-mimetic nanovesicles make them promising as drug-delivery carriers. To tap into this potential, accurate non-destructive methods to load them and characterize their contents are of utmost importance. However, their small size, polydispersity and aggregation in solution make quantitative characterizations of their loading particularly challenging. Here we develop an ad-hoc methodology based on a burst analysis of dual-color confocal fluorescence microscopy experiments, suited for quantitative characterizations of exosome-like nanovesicles and of their loading. We apply it to study bioengineered nanovesicles, loaded with dUTP cargo molecules, synthetized from detergent-resistant membranes of animal extracellular vesicles and human red blood cells. For both classes of bioengineered nanovesicles we prove, by means of dual-color fluorescence cross-correlation spectroscopy (FCCS), successful loading. Furthermore, by a dual-color coincident fluorescence burst (DC-CFB) analysis of the experimental data, we retrieve size and loading statistics for both types of nanovesicles. The procedure affords single-vesicle characterizations, which are essential for reliable quantitative studies of loading processes in exosomes and exosome-mimetic nanovesicles, especially in light of the typically high heterogeneity of their populations. Moreover, the method implementation can be easily adapted to the investigation of a variety of combinations of different cargo molecules and biological nanovesicles besides the proof-of-principle demonstrations considered in this study. The results provide a powerful characterization tool, well-suited for the optimization of loading processes of biomimetic nanovesicles and their advanced engineering for therapeutic drug delivery.

---

Exosomes are small extracellular vesicles of endocytic origin that regulate cell-to-cell communication and play a crucial role in the immune response and in the progression of several pathologies, including cancer.^1, 2^ In their outer membrane and in their lumen, they carry a wealth of biological information, in the form of proteins, lipids, and genetic material, reflecting a multitude of micro-environmental and metabolic conditions, making them extremely interesting as potential biomarkers for several diseases.^3–5^ Their small size, targeting functionalities and low immunogenicity make them also particularly promising as therapeutic drug-delivery vesicles.^6–13^ However, their large-scale deployment as drug-delivery carriers is adversely affected by the low extraction yields of exosomes naturally arising in biological fluids and by the elaborate procedures required for their purification.^14, 15^ Moreover, their use as drug carriers is further hampered by possible therapeutic crosstalk with their intrinsic biological cargo and by their relatively low drug-loading yields.^16^ The need to overcome these limitations has spurred significant research on the bioengineering of exosome-mimetic nanovesicles.^17–22^ A number of promising approaches have recently emerged for the synthesis of exosome-mimetic vesicles, which retain most of the appealing features of their naturally occurring counterparts, while ensuring high production yields and enhanced drug-loading capabilities.^22–24^ Among them, approaches for the production and loading of nanovesicles from detergent resistant membranes (DRM) of red blood cells are particularly promising, for the perspective scalability of their production and loading processes.^25, 26^

All potential applications of exosomes and exosome-mimetic nanovesicles for therapeutic drug delivery call for the development of suitable experimental and analysis techniques to accurately quantify their loading.^27–31^ Key challenges in this respect arise from the very small sizes (30 – 150 nm in diameter) and the wide heterogeneity of natural exosomes and of their biomimetic counterparts. This imposes the need for screening on a single-vesicle basis, with high spatial resolution and yet high throughputs, in order to build relevant statistics. Well-established methods for nanoparticle characterization, such as nanoparticle tracking analysis or atomic force microscopy (AFM) are unfortunately not suited for the analysis of nanovesicle contents.^32^ Optical fluorescence spectroscopy techniques provide the most promising non-destructive approach in this respect: they are well-suited for measurements in physiological solutions, afford vesicle-by-vesicle analyses with high biomolecular specificity and sensitivity,^33–35^ and enable the simultaneous retrieval of collective properties over large ensembles of molecules and nanoparticles.^36–39^

Here we demonstrate a dual-color fluorescence microscopy method suitable for accurate quantification of the loading yields of exosomes and exosome-mimetic nanovesicles filled with cargo molecules. The method overcomes measurement challenges stemming from the typical heterogeneity of the nanovesicle populations, from their natural tendency to form aggregates and from unwanted background signals in solution. To this aim, we employ a dual-color fluorescent tagging scheme encompassing the outer membrane of the nanovesicles and their cargo molecule (dUTP) and use extracellular vesicles purified from equine seminal plasma (semEV) as a reference.^40^ By means of fluorescence cross-correlation spectroscopy (FCCS) we demonstrate the successful loading with dUTP molecules of two classes of DRM nanovesicles, derived with the procedure of Ref. 41 from either semEVs displaying exosome-mimetic characteristics, or human red blood cell ghosts. By means of dual-color coincident fluorescence microscopy, we retrieve accurate estimates of dUTP-loading yields, identify vesicle sizes at which the loading efficiency is maximized, and infer average numbers of cargo molecules per nanovesicle, for both nanovesicle typologies.

## RESULTS AND DISCUSSION

### Dual-color fluorescence measurements on dUTP-loaded nanovesicles

The samples under study consisted of bioengineered nanovesicles made of detergent resistant membranes, originating from semEVs, isolated and purified from horse seminal plasma, and red blood cell ghosts, isolated and purified from human blood. Referring to their origin, the two types of nanovesicles are designated as EXO and RBC nanovesicles, respectively, throughout the rest of the paper. The sample preparation involved multiple ultracentrifugation steps, followed by loading of the vesicles with fluorescently-tagged and membrane-impermeable dUTP molecules, as detailed in Methods and described in previous work.^22, 41^ In view of the dual-color optical measurements, the cargo molecules (dUTP) were fluorescently tagged with a green-emitting dye (Alexa488) prior to loading. After loading, the outer membrane of the nanovesicles was tagged with a lipophilic dye (CellVue Claret) (**Figure 1a**). The choice of the two dyes ensured minimal spectral overlap between their emission, located in the green (Alexa488) and far-red (CellVue Claret) spectral ranges (see **S3**). Details on sample preparation are provided in Methods.

**Figure 1.**
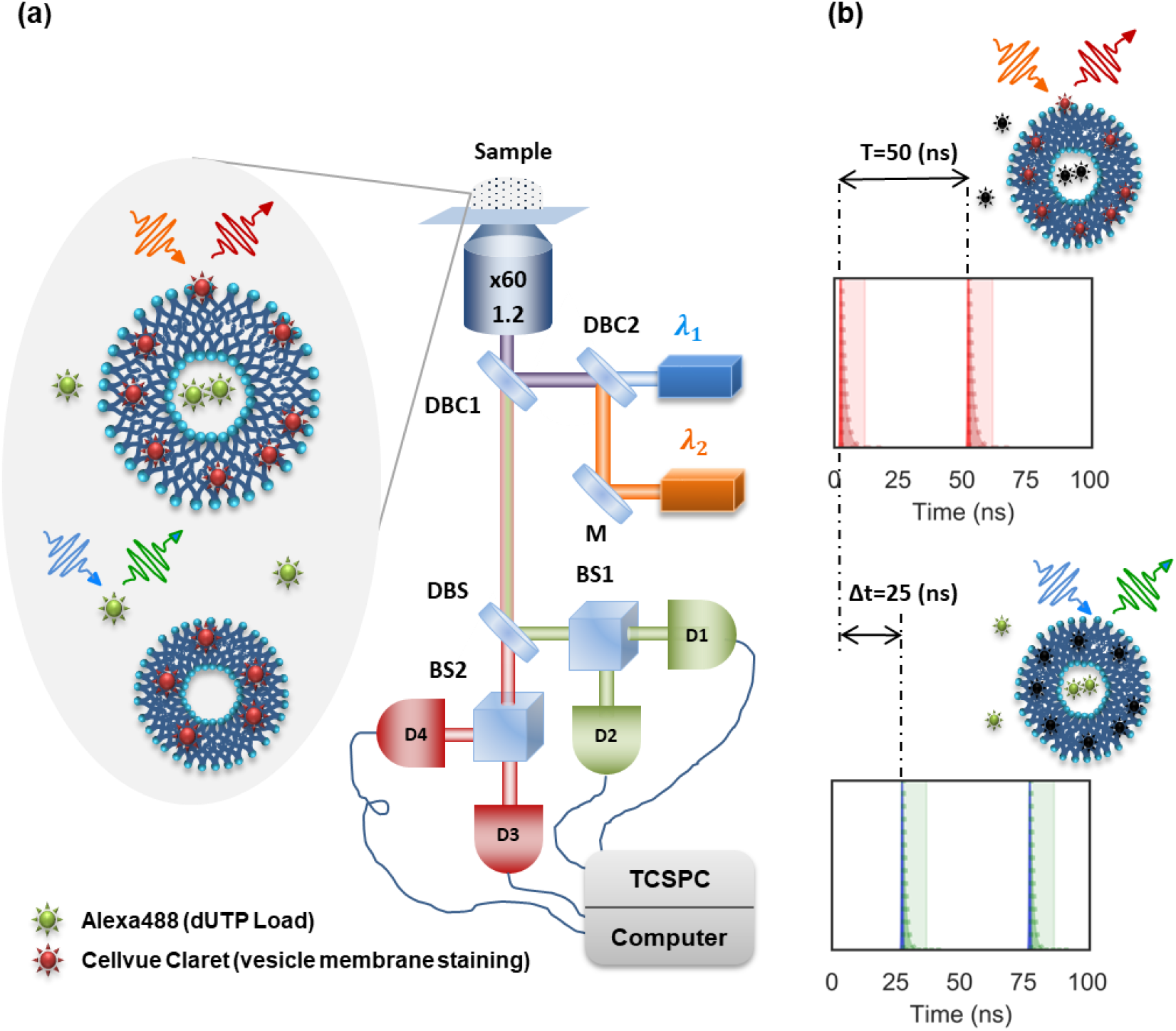
**(a)** Detail of the specimen (nanovesicle solution) and sketch of the experimental setup used for dual-color fluorescence spectroscopy with fluorescently-tagged dUTP cargo molecules (Alexa488 dye, excitation at *λ*_1_ = 485 nm, detection centered at *λ_green_* = 535 nm) and tagged nanovesicles (CellVue Claret dye, excitation at *λ*_2_ = 640 nm, detection centered at *λ_red_* = 720 nm). D1-4 = avalanche photodiodes, TCSPC = time correlated single photon counting module, DBC1-2 = dichroic beam combiners, M = mirror, BS1-2 = 50:50 beam splitters and DBS = dichroic beam splitter. **(b)** Illustration of the time-gating scheme for the red and green fluorescence time traces, implemented with 90 ps Gaussian excitation pulses (vertical lines) with a relative delay Δt =25 ns, resulting in red and green emission signals exponentially decaying over a few ns, post-selected with time-gated detection in the shaded temporal windows.

For the optical experiments, the nanovesicles were dispersed in a physiological PBS solution and analyzed with the two-color confocal microscopy setup sketched in **Figure 1a**. Two pulsed lasers provided blue (485 nm) and red (640 nm) excitations for the dyes attached to the cargo molecules (Alexa488) and the nanovesicle membrane (CellVue Claret), respectively. The emission wavelengths of the former (~675 nm) and the latter (~520 nm) were separated on the detection side by a dichroic beam splitter (DBS in **Figure 1a**) and further isolated by spectral filters centered around 720 and 535 nm (with bandwidths of 150 and 70 nm), respectively. To further improve on the discrimination between the florescence originating from the green and the red dyes, a pulse-interleaved excitation scheme was employed, with suitably synchronized post-selection windows in detection.^42, 43^ This is schematically illustrated in **Figure 1b**. The lasers emitted two pulse trains at the same repetition rate (v_rep_= 20 MHz) and interleaved in time with a temporal offset of 25 ns. The latter is significantly shorter than the nanovesicle diffusion time through the detection volume in the microscope (~ 5 ms), but longer than the emission lifetimes of the fluorophores used in the experiments (≤ 4 ns), see also **S3**.

The photon timestamps of the red and green fluorescence signals were collected by a time correlated single-photon counting module, using avalanche photodiodes. This enabled the simultaneous retrieval of separate but synchronized fluorescence intensity time-traces at the two colors and of their FCCS curves. Further details are provided in Methods and Supplementary Information (**S1**).

### Fluorescence Cross-Correlation Spectroscopy (FCCS) analysis

Fluorescence correlation spectroscopy (FCS) techniques have been successfully applied to previous exosome studies.^33, 44^ However, conventional single-color FCS methods do not lend themselves to accurate assessments of nanovesicle loading, due to the competition between the signal stemming from the latter with typically high background levels originating from residual fluorescently-tagged cargo molecules which are freely diffusing in solution. In our case, the contribution from the latter could not be completely removed from the measurements, even with additional purification procedures, hence the choice to develop a dual-color fluorescence cross correlation spectroscopy (FCCS) scheme, which encompasses tagging both the cargo molecules and the nanovesicles. Loaded nanovesicles can then be identified by cross-correlation analyses of the dual-color fluorescence signals, removing a substantial portion of the background from free dyes.

FCCS is a well-established method to study biomolecular interactions at the nanoscale.^37^ In this section we describe its adaptation to specificities of experiments on loaded exo-mimetic nanovesicles, specifically EXO and RBC loaded with dUTP cargo molecules, as shown in **Figure 2.** FCCS curves retrieved from dual-color experiments performed under identical conditions on EXO and RBC nanovesicles are shown in **Figure 2a** and **Figure 2b**, respectively. *G_red_* and *G_green_* are the measured autocorrelation curves of the fluorescence signals from the CellVue Claret and the Alexa488 dyes used to tag the nanovesicle membrane and the dUTP molecules, respectively. *G_cross_* are the cross-correlation curves of the red and green signals obtained from the same measurements.

**Figure 2.**
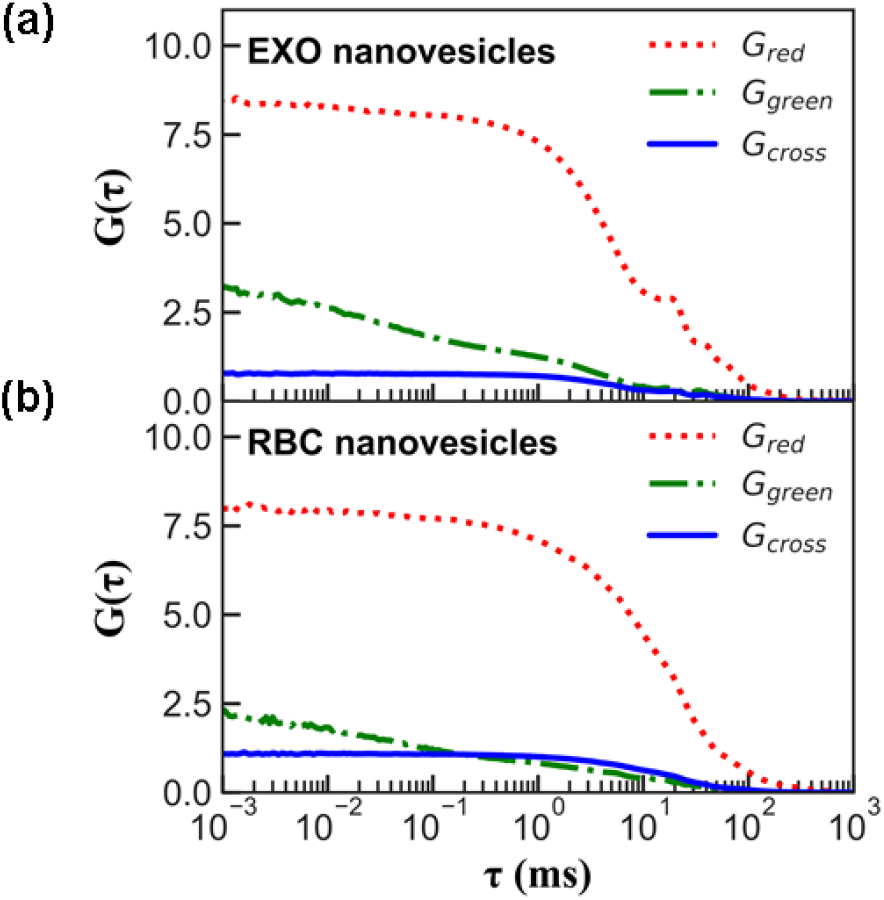
Dual-color fluorescence correlation curves recorded from: **(a)** semEV-derived and **(b)** red blood cell-derived nanovesicles. Red-dotted and green dash-dotted lines: autocorrelation curves for the fluorescence of the vesicle-membrane tag CellVue Claret (*G_red_*) and of the cargo molecule tag Alexa488 (*G_green_*), respectively. Solid blue lines: fluorescence cross-correlation curves (*G_cross_*).

The experimental conditions and the procedures used to retrieve the auto and cross correlation curves are described in Methods. Following standard FCCS protocols, numerical fits on the correlation curves allow estimates of the diffusion time and concentration of the red, green and dual-colored particles in the experiments. The results are presented in supplementary information (see **S1**). The FCCS analysis is based on the assumption of uniformly labeled particles with 1:1 binding, which is not strictly true for our samples, where there might be several and different amounts of fluorophores per vesicle. Furthermore, while FCCS is very well-suited to analyze uniform and narrow particle distributions in terms of size and brightness, its data may be more difficult to interpret for ensembles of particles with high heterogeneity and polydispersity, which is often the case for exosomes and exosome-mimetic vesicles.^45^ The FCCS results nevertheless provide useful insights on the nanovesicle populations and their loading. Specifically, the non-zero values of the cross-correlation functions *G_cross_* apparent from the data plotted in **Figure 2** clearly demonstrate the presence of loaded nanovesicles in both EXO and RBC populations. However, the retrieved *G_green_* curves appear also to deviate from a standard FCS single-particle diffusion function and are better fitted by a double-diffusion function (see also Supp. Inf.). The shortest of the two diffusion times inferred from the latter closely matches the diffusion time of free dyes. This confirms the non-negligible contribution of free dUTP molecules in solution as a limiting factor for single-color FCS measurements. The dual-color scheme overcomes this limitation and significantly improves the signal-to-noise ratio for the Alexa-stained dUTP molecules loaded in the nanovesicles, as indicated by the shape of the *G_cross_* curves (as opposed to *G_green_*), which is closer to that of a single diffusing species, with diffusion times consistent with those retrieved for *G_red_* (see also **Table S2**). The values of the diffusion times (*τ_r_* and *τ_rg_* in **Table S2**) indicate also that *G_red_* and *G_cross_* are over-biased by vesicles of larger size. In fact, the nanovesicle radii inferred by standard FCCS analysis (*R_r_* and *R_rg_* in **Table S2**) largely exceed those determined on the same nanovesicle populations through independent AFM measurements (see also following section and **Figure 3**), which indicate comparable sizes for exosomes (semEV) and exosome-mimetic nanovesicles (EXO and RBC), i.e. radii below 75 nm.^46^ The larger sizes extrapolated from FCCS measurements are consistent with the observed occurrence of nanovesicle agglomerates in solution. This effect arises naturally in physiological conditions and cannot be completely removed without disruptive chemical or mechanical treatments, which may on the other hand compromise the biological relevance of the measurements.^45, 47^ Overall the FCCS analysis demonstrated the successful loading of both EXO and RBC nanovesicles and provided estimates for their average loading yields, amounting to 12% and 22%, respectively. It also highlighted challenges in the quantification of the vesicle size stemming from the high polydispersity of these systems, from a natural tendency of vesicles to aggregate and from the presence of residual Alexa488-dUTP molecules in solution. The following sections illustrate how these issues can be resolved by a burst analysis methodology, which can be applied to the same experimental data streams underlying the FCCS results.

**Figure 3.**
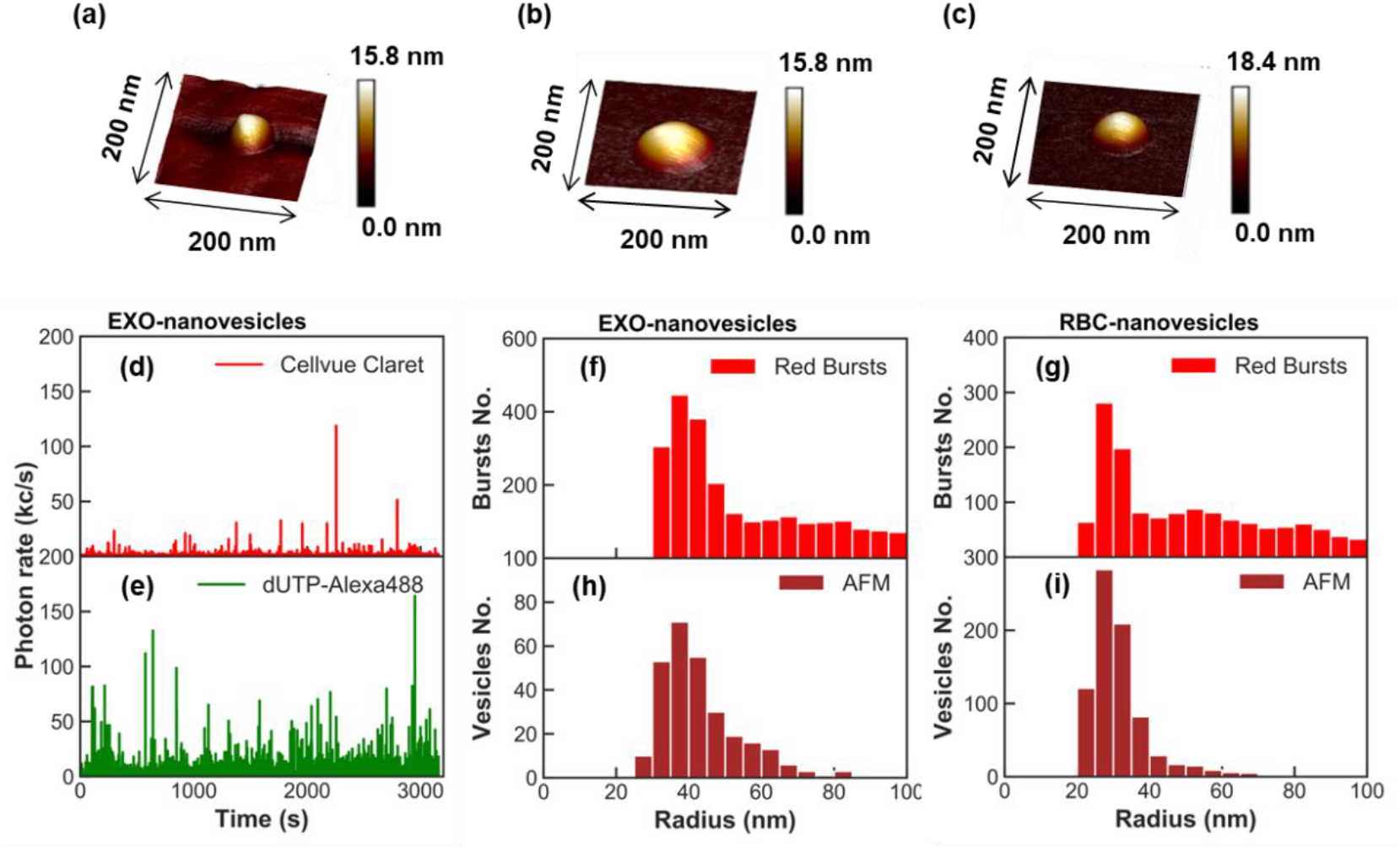
High resolution AFM images of: **(a)** semEV, **(b)** EXO and **(c)** RBC individual nanovesicles. Detail of: **(d)** red and **(e)** green fluorescence time traces from experiments on EXO. Size histograms obtained by RFB analyses for: **(f)** EXO and **(g)** RBC nanovesicles (5 nm binning). Size histograms measured by AFM for: **(h)** EXO and **(i)** RBC nanovesicles.

### Dual-Color Coincident Fluorescence Burst (DC-CFB) analysis

To extract quantitative information at a single-vesicle level from the experiments, we developed an analysis procedure starting from the time traces of the red and green signals retrieved in the dual-color experiments. Its basic idea is to identify first the bursts arising from vesicles and dUTP molecules, respectively and then analyze their coincidences in time, corresponding to dUTP-loaded nanovesicles. The procedure bears similarities with approaches used in single-molecule and Förster resonance energy transfer (FRET),^48,49,50^ and is briefly outlined in the following paragraphs.

First, to avoid crosstalk between the two colors, the photon streams at each wavelength range were time-gated with respect to the fluorescence lifetimes of the corresponding dyes, as explained in **Figure 1b**. The lifetime histograms and the time-gating windows can be found in supplementary information (**S3**). Furthermore, for a reliable burst analysis, the time-dependent background rates for each color stream were subtracted from the raw data (examples available as supplementary information, **S3**). It is important to consider the time-dependence of the background rates, especially in long measurements, due to photobleaching and possible evaporation of the solution over time. Thereafter the total number of nanovesicles was determined by counting the bursts in the red fluorescence time trace. Their size was then assessed on an individual basis from the burst duration, as described in Methods. A similar analysis was developed for the green fluorescence signal. The green and red bursts which overlap in time (dual-color coincident bursts) are used to identify the subset of loaded nanovesicles. Statistical analyses were then conducted on these bursts to extract their number and size (see Methods and **S3**). **Figure 3d(e)** shows a typical time trace for the red (green) fluorescence signal, together with the outcome of statistical analyses on such traces.

To search for the relevant bursts in the fluorescence time traces, we adapted an open-source FRET-BURST code.^42, 51^ The specificities of our DC-CFB configuration required a careful adjustment of the burst selection parameters, namely: the minimum threshold for the number of photons in each burst *M_r.g_* (where *r* and *g* correspond to red and green traces, respectively) and the minimum allowed photon count-rate *F_r.g_*, quantified as a multiple of the experimentally determined average background rate.^52^ Moreover, we introduced a specific criterion in the search for coincident bursts, concerning the relative center delay Δ*t_rg_* between time-overlapping red and green bursts. The criterion involved setting an upper threshold for Δ*t_rg_*, typically ~ 5 μs. A specific section in supplementary information (**S3**) is considers DC-CFB parameter optimization. The following section highlights key features and discusses further validation procedures adopted in the analysis.

### Fluorescence burst analysis validation by atomic force microscopy

Each burst in the red time traces (**Figure 3e**) corresponds to a vesicle diffusing through the measurement volume in the microscope setup. The outcome of the red fluorescence burst (RFB) analysis hence provides information on the total population of (loaded and unloaded) nanovesicles. The RFB results were routinely checked against independent size measurements performed by AFM on vesicles dispersed on silicon chips. Details on the AFM sample preparation, instrumental settings and image analysis procedures are given in supplementary information (**S2**). The resolution limit in such measurements is determined by the radius of the AFM tip (~8 nm), which allows individual semEV, EXO and RBC nanovesicles to be resolved equally well (**Figure 3a-c**), confirming their rounded shape and integrity in all cases. In **Figure 3f-g** we compare the size distribution obtained by RFB and AFM analyses performed on the same nanovesicle populations considered in **Figure 2**. The size distributions for EXO and RBC nanovesicles peak at radii 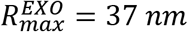 and 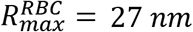, respectively, with very good agreement between the AFM and RBC results also at smaller sizes. However, at larger sizes the RFB distributions feature an increasing discrepancy with the AFM curves, predominantly due to nanovesicle aggregation in solution. Further quantitative comparisons are provided in **Figure S10** and **Table S3**. In light of these observations and consistently with the size range attributed to extracellular vesicles,^46^ we therefore set an upper limit to the nanovesicle size range considered reliable for statistical RFB analyses, equal to a radius of 75 nm. This yields average nanovesicles radii 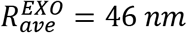 and 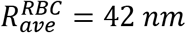, which compare favorably with the AFM results (see also **Table S3**). In the next sections we show how reliable quantitative assessment of the loading yields can also be retrieved from the experimental data by a suitable coincidence burst analysis methodology.

### Dual-Color Coincident Fluorescence Burst (DC-CFB) analysis

Coincident bursts in the red and green fluorescence time traces identify the sub-population of dUTP-loaded vesicles. The key parameters for the DC-CFB analysis are the thresholds for the photon number, the count rates in the red and green bursts and the time tolerance window for identification of their temporal coincidence (*M_r_, F_r_, M_g_, F_g_* and Δ*t_rg_*). The correct settings of *M_r_* and *F_r_* were verified by comparing the results of the RFB size distributions with independent AFM measurements, as previously discussed. The optimal values of *M_g_* and *F_g_* were instead determined with an ad-hoc optimization methodology based on physical insights specific to the system under study. Essentially, we used the fact that the dUTP-loaded nanovesicles can be identified in two ways, namely: 1) as coincident bursts singled out from the red time trace, which we shall refer to as *coincident-red* bursts, or 2) as coincident bursts singled out from the green trace, referred to as *coincident-green* bursts. Accordingly, two size distributions can be extracted from the burst analysis of dual-color experimental data streams. They are shown in **Figure 4** for the same vesicle populations considered in **Figure 2a-b** and **Figure 3f-g**. *Coincident-red* traces for EXO and RBC nanovesicles are shown in **Figure 4a-b**. The corresponding *coincident-green* traces are shown in **Figure 4c-d**.

**Figure 4.**
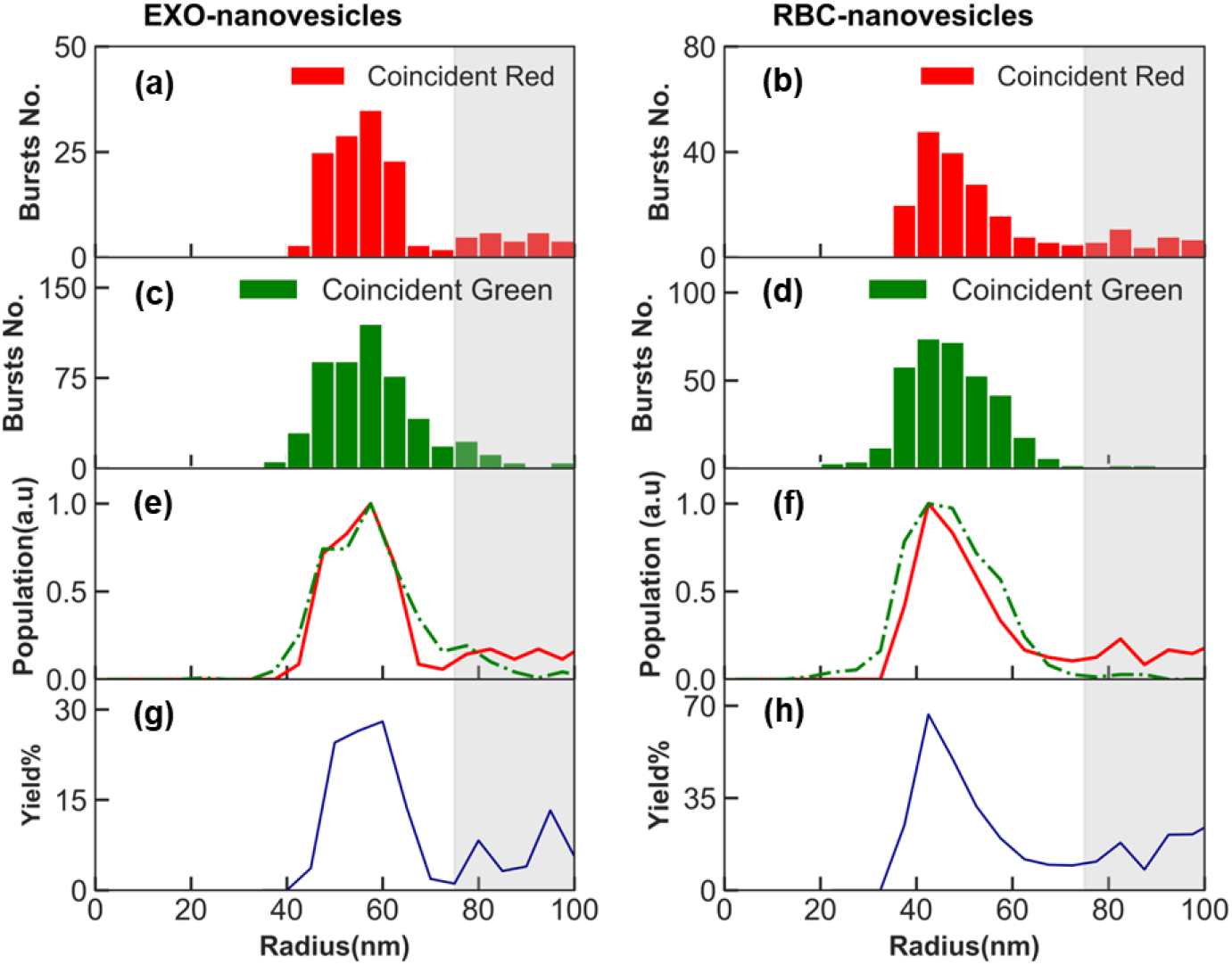
Dual color coincident fluorescence burst analysis with optimized parameter settings (see also Supplementary Information **S3**). Size histograms for: **a-b**) *coincident-red* and **c-d**) *coincident-green* bursts. **e-f)** Normalized *coincident-red* (red solid) and *green* (dash-dotted green) distributions and **g-h)** loading yields as a function of nanovesicle radius. EXO (left) and RBC (right column). The grey-shading marks the region *R* > 75 nm, beyond the confidence range for single-vesicle analyses due to aggregation effects (see also discussion in the text).

In an ideal case, the size histograms inferred from the two coincident burst analyses should be identical, as they record individual occurrences of the same physical event, i.e. the diffusion of a dUTP-loaded nanovesicle in the detection volume. However, depending on the parameter settings, the distributions do vary. Minimizing the difference between *coincident-red* and *coincident-green* size distributions proved to be a viable and robust criterion to find the optimal settings for the DC-CFB analysis across all measurements. Moreover, at the excitation power used in our experiments (80 *μ*W at most, corresponding to ~40 and 25 kW/cm^2^ for the green and red channels, respectively), we could rule out a significant impact of fluorophore saturation and photobleaching effects on the results.^53, 54^ Specifically, for the green traces we evaluated a possibility for only 7% (5%) of the EXO (RBC) loaded vesicles at the peak radius size of 60 nm (42 nm) to be bleached during diffusion through the excitation spot, in light of reported photostability measurements on the Alexa488 fluorophore.^55^ As for the red burst analysis, the impact of photobleaching can be expected to be even lower. Further details on the DC-CFB analysis settings and photobleaching estimates are provided in Supporting Information (**Figures S11-S15** and **Table S4**). **Figure 4e** and **4f** show the final results, illustrating the level of agreement attained between the *concident-red* and *coincident-green* distributions through an optimal choice of the DC-CFB parameters, determined by minimizing the target error function in the size range typical for exosome-like nanoparticles (*R* < 75 nm). Even if the difference between the red and green distributions in **Figure 4e-f** increases in the grey-shaded regions (*R* > 75 nm), a very good agreement is obtained around the peak, located at a radius of 60 *nm* and 42 *nm* for EXO and RBC nanovesicles, respectively. Such values are higher than those obtained by the RFB analysis (37 and 27 *nm*, respectively) on the overall nanovesicle populations, which indicates a slightly larger size of the most densely populated dUTP-loaded nanovesicles with respect to unloaded ones (**Table S3** and **Table II**). This is a feature we consistently observed across all measurements and nanovesicle typologies we have investigated so far.

**Table II.** DC-CFB analysis results for EXO and RBC nanovesicles. 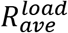 is the average nanovesicle size (radius), *Yield_ave_* the average loading yield, 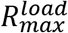 the predominant size of loaded nanovesicles and *Yield_max_* the loading yield for 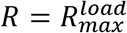. (*N_cargo_/vesicle*)_*ave*_ is the extrapolated average number of dUTP cargo molecules per nanovesicle.

| Sample | $R_{ave}^{load}$ | $Yield_{ave}$ | $R_{max}^{load}$ | $Yield_{max}$ | $\left(\frac{N_{cargo}}{vesicle}\right)_{ave}$ |
| --- | --- | --- | --- | --- | --- |
| **EXO nanovesicles** | 55±6 nm | 12% | 60±5 nm | 28% | 1.50 |
| **RBC nanovesicles** | 49±9nm | 20% | 42±5 nm | 67% | 1.15 |

### Loading yields

The single-vesicle tracking capability of the DC-CFB approach allows the loading yield to be estimated as a function of nanovesicle radius: 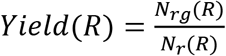, where *N_rg_*(*R*) is the number of coincident red bursts evaluated by the optimized DC-CFB procedure described previously and *N_r_*(*R*) is the total number of red bursts obtained from the single-color RFB analysis. **Figure 4g-h** plot the loading yield as a function of EXO and RBC vesicle sizes. In both cases, the most frequent size of the loaded exosome-mimetic nanovesicle sub-population is slightly larger than that of the overall nanovesicle population (**Figure 4a-b**). From the data shown in **Figure 4g-h**, a maximum loading efficiency *Yield_max_* equal to 28% and 67% is inferred for EXO and RBC nanovesicles, respectively. However, a much more meaningful figure of merit is given by the value of the loading yield averaged over the whole size distribution and defined as: 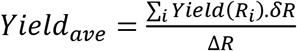, where Δ*R* is the size range at which the yield is non-zero and *δR* is the binning value of the histograms (*δR* = 5 nm, see also Supp. Inf. **S3**). The average yield was also found to be consistent across repeated measurements on the same vesicle populations at different times and independent of the binning values (in the range 5-15 nm) chosen in the analysis. The values of the average loading yields estimated from the DC-CFB analysis are 12% for EXO and 20% for RBC nanovesicles, in excellent agreement with the outcomes of the FCCS analysis (see also **Table S2**).

Furthermore, the data retrieved by the DC-CFB method can be used to estimate the average number of dUTP molecules loaded in each nanovesicle. This is done under the assumption of uniform brightness of the green fluorescence signal originating from the cargo molecules and following calibrations of the average photon count rate of single dUTP-Alexa488 molecules in solution (performed with the same experimental settings in the setup of **Figure 1**). One can then normalize the photon count rate of each burst in the *coincident-green* signal to the average photon count rate of the dUTP-dye molecule to infer the approximate number of cargo molecules per vesicle. These data can be plotted together with the information derived from the temporal duration of the bursts providing the size of each loaded nanovesicle. The outcome of the analysis is displayed in the two-dimensional histograms of **Figure 5**, showing the 2D distribution (as a color map) of the loaded nanovesicles as a function of vesicle size and number of cargo molecules per vesicle (see also Methods). The 2D map indicates the number of cargo molecules per carrier vesicle and the size of the latter. The distribution has a peak at approximately 1.1 (1.0) number of cargo molecules per vesicle and at a radius size of 60 nm (42 nm) for EXO (RBC) samples, as highlighted by the darker spots in the 2D maps. The horizontal and vertical histograms in the same pictures show the distributions in size and number of cargo molecules of the loaded nanovesicle subpopulations, retrieved from the *coincident-red* and *coincident-green* bursts, respectively. From the vertical histogram the average number of loaded cargo molecules per nanovesicle 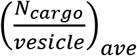 is estimated to be 1.50 and 1.15 for EXO and RBC nanovesicles, respectively.

**Figure 5.**
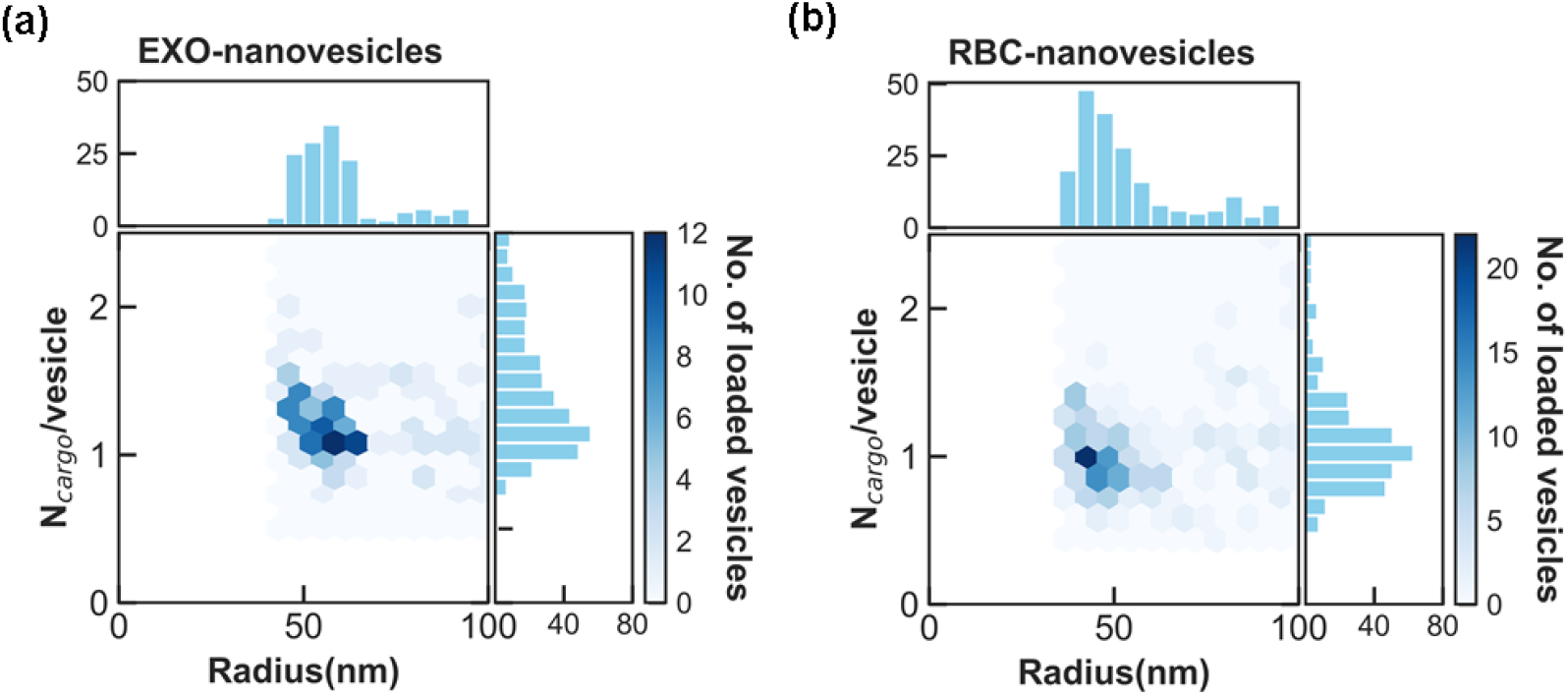
Loaded nanovesicle subpopulation analysis. **a)** Two dimensional histograms of loaded EXO nanovesicles versus vesicle size and number of loaded cargo molecules, evaluated from *coincident-red* (CellVue Claret) and *coincident-green* (Alexa488) burst analyses, respectively. The size and number of cargo molecules per vesicle for the loaded vesicles are depicted in the horizontal and vertical subplots, respectively. **b)** Same as **a)** for loaded RBC nanovesicles.

**Table II** summarizes the results of the full DC-CFB analysis for dUTP-loaded EXO and RBC samples.

In conclusion, we have established a dual-color coincident fluorescence burst (DC-CFB) methodology for quantitative non-destructive studies of heterogeneous populations of fluorescently-tagged exosome-mimetic nanovesicles and their cargos. The technique has singlemolecule sensitivity and allows characterizations on a vesicle-by-vesicle basis. Given the typically high heterogeneity of biological nanovesicle populations, these enhanced capabilities are essential for reliable assessments of collective properties, such as loading yields with cargo molecules for therapeutic purposes. We applied the DC-CFB approach to evaluate the efficiency of the loading process of dUTP cargo molecules into nanovesicles synthesized from horse prostasomes, i.e. extracellular vesicles derived from seminal plasma (EXO), and from human erythrocytes (RBC). Both studies confirmed the reliability and accuracy of the method in spite of measurement challenges associated with the presence of residual free dyes, high polydispersity and aggregation of the biological nanovesicles in solution.

The outcomes of single-color fluorescence burst analyses were also validated against independent AFM measurements. Good agreement was found between the two methods, apart from a discrepancy for larger vesicle sizes, induced by their tendency to form aggregates in solution. An empirical criterion was introduced to calibrate and optimize the coincident bursts analysis of the dual-color photon streams stemming from loaded nanovesicles. The method was then applied to size-resolved quantitative evaluations of the loading efficiencies of EXO and RBC nanovesicles with dUTP cargo molecules. Quantitative evaluations of the average dUTP-loading yields obtained by the DC-CFB analysis for the two nanovesicle typologies considered in the study yielded values of 12% for EXO and 20% for RBC specimens, in excellent agreement with FCCS measurements, providing values of 12% and 22%, respectively. Moreover, the DC-CFB analysis indicated also that loaded vesicles are larger, on average, than the overall population of (loaded and unloaded) nanovesicles they belong to.

The results demonstrate a new approach for the study of bioengineered exo-mimetic nanovesicles and of their cargo. The method is well-suited for quantitative analyses and provides reliable assessments of loading efficiencies. It can reliably track nanovesicle properties as a function of different preparation conditions and nanovesicle/cargo combinations, hence providing a novel and powerful tool to drive further progress in the use of exo-mimetic nanovesicles for therapeutic purposes. Extension of the method to the study of autofluorescent molecules in native exosome cargos, as opposed to synthetic drug-loaded nanovesicles, may also pave the way to a deployment of this approach for diagnostic applications in exosome-based liquid biopsy.

## METHODS

### Preparation of semEV (extracellular vesicles from stallion semen)

Pooled seminal plasma from stallions was centrifuged for 30 min at 10,000g and 4°C (Beckman Coulter, BC - rotor SW32Ti) to pellet cell debris and larger molecular complexes. The supernatant was saved and transferred to a new tube. Supernatants were ultracentrifuged for 1h at 100,000g and 4°C (BC-SW32Ti) to pellet vesicles. Saved vesicle-pellets were resuspended in phosphate buffered saline (PBS) and separated in a density gradient built with 50%, 42.5% (1.19 g/cm^3^), 30% (1.13 g/cm^3^), 15% sucrose (with the sample top-loaded) for 4h at 160,000g and 4°C (BC-SW40Ti). Seminal fluid extracellular vesicles that were layered on 40% sucrose (density range 1.13-1.19 g/cm^3^), hereby denoted as semEV, were suspended in PBS and pelleted by ultracentrifugation for 1h at 100,000g and 4°C (BC-SW32Ti). Pellets were resuspended in PBS and semEV concentrations were estimated by BSA protein kit (Thermo Fisher, Pierce BCA Protein Assay Kit) and adjusted to 2mg/mL. Prepared semEV were frozen at −20°C until use.

### Preparation of red blood cell ghosts

Blood bags containing red blood cells were purchased from Uppsala University Hospital. Blood donors had been informed and agreed upon use in scientific research on donated blood. All blood bags had been deidentified. Ten mL of red blood cellswere washed 3 times in PBS (1:5) by centrifugation for 10 min at 2100g and 4°C (Nino lab, Heraeus Multifuge3s). Washed red blood cells were lysed in hypotonic phosphate buffer (PB, 53.4 mOsmol/L) and repeatedly washed in PB until most of the hemoglobin was removed (5-8 times with RBC:PB = 1:4) by ultracentrifugation for 30 min at 20,000g and 4°C (BC-SW32Ti). Washed red blood cell ghosts were kept at −20°C for less than three months.

### Preparation and loading of nanovesicles

Stored semEV (8 mg, estimated by BCA protein kit) and stored red blood cell ghosts (10mL, exact estimation of protein amount was impracticable because of remaining hemoglobin molecules) were suspended in PBS and ultracentrifuged for 1h at 100,000g and 4°C (BC-SW32Ti). Pellets were resuspended in PBS and mixed with Triton X100 (detergent) in a final concentration of 1% detergent followed by incubation for 30 min on ice. The detergent treated samples were separated on density gradient built with 50%, 30%, 24%, 10% sucrose (with the sample on top) by another ultracentrifugation for 5h at 230,000g and 4°C (BC-SW40Ti). Fraction with buoyancy at 30% sucrose (1.13g/cm^3^), corresponding to detergent-resistant membrane (DRM) vesicles, was collected and pelleted by ultracentrifugation in PBS containing 10% sucrose for 1h at 150,000g and 4°C (BC-SW32Ti). Pellets were kept at −20°C until use.

Loading was performed by adding Alexa488-dUTP (Thermo Fisher, ChromaTide Alexa Fluor 488-5-dUTP, a single nucleotide) diluted (1:50) in PBS, directly on the frozen DRM vesicle-pellets. The physiological buffer induced a transition from hypertonicity (caused by sucrose) to isotonicity (caused by PBS), a phenomenon called “post-hypertonic lysis”.^56^ This implies an osmotic lysis of the vesicles, with rupture and revesiculation, whereby the restored vesicles supposedly incorporate the surrounding dUTP molecules in PBS, which do not display a free permeability across biological membranes. The dUTP-loaded nanovesicles were henceforth protected from direct exposure to light. They were pelleted by ultracentrifugation for 1h at 100,000g and 4°C (BC-SW32Ti). This new pellet was resuspended in CellVue Claret membrane kit-staining-component Diluent C (Sigma-Aldrich, Miniclaret-1kt). Staining of membranes took place according to manufacturer’s instructions and was stopped by adding 2% bovine serum albumin. The stained samples were floated in a density gradient built with 40%, 30%, 10% sucrose with stained sample on top, by ultracentrifugation for 5h at 230,000g and 4°C (BC-SW40Ti).

The fraction at density 1.13g/cm^3^ (30% sucrose) containing purified, loaded and stained, nanovesicles was collected and pelleted by ultracentrifugation for 1h at 100,000g and 4°C (BC-SW32Ti). Pellets (“EXO” originating from semEV and “RBC” originating from erythrocyte ghosts) were resuspended in PBS and kept at 4°C in dark until analyses.

### FCCS Setup

The schematic of the experimental setup for dual-color Fluorescent Cross Correlation Spectroscopy (FCCS) is depicted in **Figure 1a**. It consists of a commercial, epi-illuminated, confocal laser scanning microscope (Olympus FV1200) equipped with a water immersion objective (60x, NA 1.2, Olympus, UPlanSApo) and the pinhole set to 50*μ*m. Excitation sources were Picoquant lasers (LDH-D-C-485 laser and LDH-D-C-640), either running separately or in pulsed-interleaved mode (PIE), each laser running at 20MHz and with a power of 80*μ*W at the back-focal plane of the objective for all measurements. The experimental setup for FCCS measurements used detection sets made of two avalanche photodiodes for each wavelength. The fluorescence light was separated from the excitation light by a ZT405/488/635rpc-UF2 dichroic mirror (Chroma) followed by another dichroic mirror separating the green and red emission, to allow for cross-correlation between the signals from the two fluorophores. The red emission was collected through a HQ720/150 (Chroma) filter and green emission through a HQ535/70 (Chroma) filter. The emitted fluorescent light at each wavelength was split by a 50:50 beam-splitter and focused onto either Picoquant tau-spad or Perkin & Elmer (SPCM-AQR-14) detectors, all connected to a TCSPC module (HydraHarp 400). The Symphotime software (Picoquant) was used for data acquisition. Diffusion-time and counts per molecule of Rhodamine 110 and Cy5 were used as references on each day of measurements to confirm the consistency of the measurement settings. The focal volumes were determined by measuring the correlation-curve of Rhodamine110 and Cy5, with known diffusion coefficients, and fitting the experimental curves with single-diffusion profiles.

### Dual-color fluorescence measurements

In each measurement, the setup was calibrated initially with free dUTP-Alexa488 molecules in solution, to estimate the photon count rate of each green dye. Nanovesicle samples taken from a refrigerator were gently spun in a bench centrifuge for 2 minutes, before each dual-color measurement. Following 1:100 dilutions in PBS, a volume of 100*μ*L of the solution sample (~2*μ*g) was put on a glass slide on top of the microscope objective. All the fluorescence measurements were done at a constant temperature and in a completely dark room. To obtain reliable statistics on the different sizes of loaded nanovesicles, repeated measurements were made, on both types of samples (EXO and RBC) over time durations of ~50 minutes.

### FCS and FCCS analysis

Autocorrelation curves for red and green signals were evaluated from the measured time evolution of their fluorescence intensity *F*(*t*), according to:

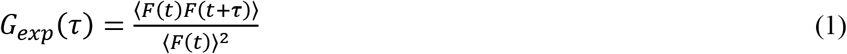

where *τ* denotes the correlation time and the angle brackets represent time average. The crosscorrelation curves were evaluated according to the following equation:

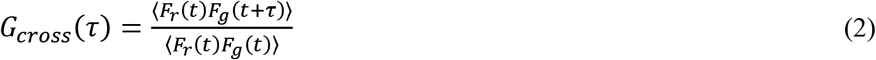

where *F_r_*(*t*) and *F_g_*(*t*) represent the evolution of the red and greed fluorescence signals in time. Details on the fits of the experimental correlation curves with standard FCS and FCCS profiles are provided in supplementary information (**S1**).

### AFM measurements

A commercial FastScan Bruker system was used to visualize the topography of vesicles at high resolution and slow scan rates (0.5 Hz) by atomic force microscopy. All measurements were performed in tapping mode in air using NCHV-A (Bruker) cantilevers made of antimony n-doped silicon. The highest possible resolution of the two-dimensional AFM images is around 8 nm which is equal to the radius of the applied tip. The size distributions of all the vesicles were quantified statistically, by taking bigger AFM images over a scan area of (5 μm)^2^ for each sample. To retrieve the overall size distribution of the nanovesicles, the two-dimensional AFM images were processed with imaging recognition routines in Matlab, to detect individual nanovesicles and fit them to circular shapes. The substrates for the AFM experiments were ~(1 cm)^2^ chips, cleaved from commercial silicon wafers (p-type). The chips were rinsed in acetone and then isopropanol, using mild sonication for 5 minutes in each solution, and then dried using a nitrogen gas flow. Prior to deposition on chip, the vesicle samples were diluted by (1:10) dilution series in double-distilled water, up to 10 times, to allow good dispersal of the vesicles on the silicon chip surface. Dilutions of both vesicle types were spun using a bench centrifuge. A drop of 1μL from each dilution was pipetted and dispensed manually on silicon chips and then allowed to dry for 3 hours at ambient temperature.

### DC-CFB analysis

For the dual-color coincident fluorescence burst analysis, the photon timestamps (50 ns time binning) in the red and green channels were first time-gated according to their excitation pulses and corresponding fluorescence dye lifetimes (**Figure S9**). Before the burst search, the time-dependent background rates for the red and green channels were evaluated (**Figure S10**) and subtracted from the corresponding photon time-stamps. The parameters utilized for the burst search in the red and green channels under optimized conditions are listed in **Table S4**. The radius (*R*) of the nanovesicles was assessed on an individual basis from the burst duration time (*τ_burst_*) according to Stokes–Einstein diffusion theory, i.e. through the formula:

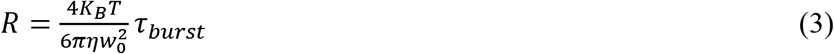

in which *K_B_*, *T, η* are the Boltzmann constant, the lab temperature and the viscosity of the solvent (water), respectively, while *w*_0_ is the lateral radius of the confocal volume. Temporally overlapping red and green bursts were identified as coincident with a tolerance Δ*t_rg_*, as discussed in the text (see also **Figure S11**). The optimal settings of the DC-CFB analysis were determined by comparing the normalized *coincident-red* and *coincident-green* size histograms in each measurement (**Figure S4e-f**) and minimizing their difference (taken as the error signal for parameter optimization). The loading yield was evaluated as the ratio of the coincident burst count over the total red burst count, typically using histograms with 5 nm binning. More details are given in supplementary information (**S3**). Moreover, to estimate the count rate of loaded nanovesicles, the photon count of green coincident bursts was divided by their corresponding time width. Therefore, the average brightness of loaded nanovesicles was calculated by dividing the count rate of coincident-green bursts by the average count rate of single green cargo (≅9.5 kcounts/s). The average count rate of dUTP.Alexa488 was measured at the same excitation power (80*μ*W) and during similar measurement time at 1 nM concentration.

## ASSOCIATED CONTENT

### Supporting Information available

Supporting information is available free of charge at http://pubs.acs.org and includes details on the theory and calibration of the FCCS experiments, AFM image analysis, comparative AFM and red burst analyses, DC-CFB analysis implementation and validation.

## Author Contributions

K. G. and M. S. conceived the study. K. G. R. was responsible for the parts of the study that involved identity, biology, design and loading of the nanovesicles from two different sources. K. G. R. and J. M. M. prepared the samples. M. S. performed the AFM measurements and their analysis. J.W. and E.S. developed the FCCS setup. M. S. and E. S. performed the fluorescence measurements. M. S. and K. G. developed the DC-CFB data analysis and wrote the manuscript with contributions and approval from all authors.

## Competing interests

K. G. R. is employed by Oblique Therapeutics AB.

## Funding Sources

The work was supported by the program for biological pharmaceutics of the Swedish Innovation Agency (Vinnova grant no 2017-02999), and by the OQS Research Environment for Optical Quantum Sensing of the Swedish Research Council (VR grant no 2016-06122). K.G. gratefully acknowledges further support from the Knut and Alice Wallenberg Foundation through the Wallenberg Center for Quantum Technology (WACQT) and from the Swedish Research Council via grant VR 2018-04487.

## Acknowledgement

We thank Owe Orwar and Carolina Trkulja from Oblique Therapeutic AB for valuable feedback and discussions, the Blood-donors organization in Uppsala for the blood samples, and Gunilla Martinsson from the National Stud, Celle, Germany, for providing the stallion seminal plasma used in the preparation of semEV.

## ABBREVIATIONS

AFM: atomic force microscopy
DC-CFB: dual-color coincident fluorescence bursts
DRM: Detergent Resistant Membranes
semEV: extracellular vesicles purified from equine seminal plasma
EXO: detergent resistant membrane vesicles derived from semEV
RBC: detergent resistant membrane vesicles derived from red blood cell ghosts
FCCS: fluorescence cross correlation spectroscopy
TCSPC: time correlated single photon counting

## Notes

### Competing Interest Statement

The authors have declared no competing interest.

## REFERENCES

1. Raposo, G.; Stoorvogel, W., Extracellular vesicles: Exosomes, microvesicles, and friends. Journal of Cell Biology 2013, 200 (4), 373–383.

2. Zhang, Y.; Liu, Y.; Liu, H.; Tang, W. H., Exosomes: biogenesis, biologic function and clinical potential. Cell Biosci 2019, 9, 19–19.

3. Lee, W.; Nanou, A.; Rikkert, L.; Coumans, F. A. W.; Otto, C.; Terstappen, L. W. M. M.; Offerhaus, H. L., Label-Free Prostate Cancer Detection by Characterization of Extracellular Vesicles Using Raman Spectroscopy. Analytical Chemistry 2018, 90 (19), 11290–11296.

4. Zlotogorski-Hurvitz, A.; Dekel, B. Z.; Malonek, D.; Yahalom, R.; Vered, M., FTIR-based spectrum of salivary exosomes coupled with computational-aided discriminating analysis in the diagnosis of oral cancer. Journal of Cancer Research and Clinical Oncology 2019, 145 (3), 685–694.

5. Cheng, M.; Yang, J.; Zhao, X.; Zhang, E.; Zeng, Q.; Yu, Y.; Yang, L.; Wu, B.; Yi, G.; Mao, X.; Huang, K.; Dong, N.; Xie, M.; Limdi, N. A.; Prabhu, S. D.; Zhang, J.; Qin, G., Circulating myocardial microRNAs from infarcted hearts are carried in exosomes and mobilise bone marrow progenitor cells. Nature Communications 2019, 10 (1), 959.

6. Pullan, J. E.; Confeld, M. I.; Osborn, J. K.; Kim, J.; Sarkar, K.; Mallik, S., Exosomes as Drug Carriers for Cancer Therapy. Molecular Pharmaceutics 2019, 16 (5), 1789–1798.

7. Snell, A. A.; Neupane, K. R.; McCorkle, J. R.; Fu, X.; Moonschi, F. H.; Caudill, E. B.; Kolesar, J.; Richards, C. I., Cell-Derived Vesicles for in Vitro and in Vivo Targeted Therapeutic Delivery. Acs Omega 2019, 4 (7), 12657–12664.

8. de Jong, O. G.; Kooijmans, S. A. A.; Murphy, D. E.; Jiang, L.; Evers, M. J. W.; Sluijter, J. P. G.; Vader, P.; Schiffelers, R. M., Drug Delivery with Extracellular Vesicles: From Imagination to Innovation. Accounts of Chemical Research 2019, 52 (7), 1761–1770.

9. Li, Z.; Zhou, X.; Wei, M.; Gao, X.; Zhao, L.; Shi, R.; Sun, W.; Duan, Y.; Yang, G.; Yuan, L., In Vitro and in Vivo RNA Inhibition by CD9-HuR Functionalized Exosomes Encapsulated with miRNA or CRISPR/dCas9. Nano Letters 2019, 19 (1), 19–28.

10. El Andaloussi, S.; Mäger, I.; Breakefield, X. O.; Wood, M. J. A., Extracellular Vesicles: Biology and Emerging Therapeutic Opportunities. Nat. Rev. Drug Discovery 2013, 12, 347.

11. Tatischeff, I.; Alfsen, A., A New Biological Strategy for Drug Delivery: Eucaryotic Cell-Derived Nanovesicles. J. Biomater. Nanobiotechnol. 2011, 2, 494.

12. Vader, P.; Mol, E. A.; Pasterkamp, G.; Schiffelers, R. M., Extracellular vesicles for drug delivery. Adv Drug Deliver Rev 2016, 106, 148–156.

13. Deshmukh, S. K.; Khan, M. A.; Singh, S.; Singh, A. P., Extracellular Nanovesicles: From Intercellular Messengers to Efficient Drug Delivery Systems. ACS Omega 2021, 6 (3), 1773–1779.

14. Cheng, Y.; Qu, X.; Dong, Z.; Zeng, Q.; Ma, X.; Jia, Y.; Li, R.; Jiang, X.; Williams, C.; Wang, T.; Xia, W., Comparison of serum exosome isolation methods on co-precipitated free microRNAs. PeerJ 2020, 8, e9434–e9434.

15. Ludwig, N.; Whiteside, T. L.; Reichert, T. E., Challenges in Exosome Isolation and Analysis in Health and Disease. Int J Mol Sci 2019, 20 (19).

16. Donoso-Quezada, J.; Ayala-Mar, S.; González-Valdez, J., State-of-the-art exosome loading and functionalization techniques for enhanced therapeutics: a review. Crit Rev Biotechnol 2020, 40 (6), 804–820.

17. Antimisiaris, S. G.; Mourtas, S.; Marazioti, A., Exosomes and Exosome-Inspired Vesicles for Targeted Drug Delivery. Pharmaceutics 2018, 10 (4).

18. Kooijmans, S. A. A.; Vader, P.; van Dommelen, S. M.; van Solinge, W. W.; Schiffelers, R. M., Exosome mimetics: a novel class of drug delivery systems. Int J Nanomed 2012, 7, 1525–1541.

19. Vader, P.; Schiffelers, R. M., ADDR editorial “Biologically-inspired drug delivery systems”. Adv Drug Deliver Rev 2016, 106, 1–2.

20. Garcia-Manrique, P.; Matos, M.; Gutierrez, G.; Pazos, C.; Blanco-Lopez, M. C., Therapeutic biomaterials based on extracellular vesicles: classification of bio-engineering and mimetic preparation routes. Journal of Extracellular Vesicles 2018, 7 (1).

21. Goh, W. J.; Zou, S.; Lee, C. K.; Ou, Y.-H.; Wang, J.-W.; Czarny, B.; Pastorin, G., EXOPLEXs: Chimeric Drug Delivery Platform from the Fusion of Cell-Derived Nanovesicles and Liposomes. Biomacromolecules 2018, 19 (1), 22–30.

22. Dubois, L.; Löf, L.; Larsson, A.; Hultenby, K.; Waldenström, A.; Kamali-Moghaddam, M.; Ronquist, G.; Ronquist, K. G., Human erythrocyte-derived nanovesicles can readily be loaded with doxorubicin and act as anticancer agents. Cancer Research Frontiers 2018, 4 (1), 13–26.

23. Xiong, F.; Ling, X.; Chen, X.; Chen, J.; Tan, J.; Cao, W.; Ge, L.; Ma, M.; Wu, J., Pursuing Specific Chemotherapy of Orthotopic Breast Cancer with Lung Metastasis from Docking Nanoparticles Driven by Bioinspired Exosomes. Nano Letters 2019, 19 (5), 3256–3266.

24. Usman, W. M.; Pham, T. C.; Kwok, Y. Y.; Vu, L. T.; Ma, V.; Peng, B. Y.; San Chan, Y.; Wei, L. K.; Chin, S. M.; Azad, A.; He, A. B. L.; Leung, A. Y. H.; Yang, M. S.; Shyh-Chang, N.; Cho, W. C.; Shi, J. H.; Le, M. T. N., Efficient RNA drug delivery using red blood cell extracellular vesicles. Nature Communications 2018, 9.

25. Li, S.-p.; Lin, Z.-x.; Jiang, X.-y.; Yu, X.-y., Exosomal cargo-loading and synthetic exosome-mimics as potential therapeutic tools. Acta Pharmacologica Sinica 2018, 39 (4), 542–551.

26. Malhotra, S.; Dumoga, S.; Sirohi, P.; Singh, N., Red Blood Cells-Derived Vesicles for Delivery of Lipophilic Drug Camptothecin. ACS Appl Mater Interfaces 2019, 11 (25), 22141–22151.

27. He, C.; Zheng, S.; Luo, Y.; Wang, B., Exosome Theranostics: Biology and Translational Medicine. Theranostics 2018, 8 (1), 237–255.

28. Shao, H. L.; Im, H.; Castro, C. M.; Breakefield, X.; Weissleder, R.; Lee, H. H., New Technologies for Analysis of Extracellular Vesicles. Chem Rev 2018, 118 (4), 1917–1950.

29. Chiriacò, M. S.; Bianco, M.; Nigro, A.; Primiceri, E.; Ferrara, F.; Romano, A.; Quattrini, A.; Furlan, R.; Arima, V.; Maruccio, G., Lab-on-Chip for Exosomes and Microvesicles Detection and Characterization. Sensors (Basel) 2018, 18 (10), 3175.

30. van der Pol, E.; Hoekstra, A. G.; Sturk, A.; Otto, C.; van Leeuwen, T. G.; Niewland, R., Optical and non-optical methods for detection and characterization of microparticles and exosomes. Journal of Thrombosis and Haemostasis 2010, 8 (12), 2596–2607.

31. Shpacovitch, V.; Hergenröder, R., Optical and surface plasmonic approaches to characterize extracellular vesicles. A review. Analytica Chimica Acta 2018, 1005, 1–15.

32. Cressey, D., Tiny traits cause big headaches. Nature 2010, 467 (7313), 264–265.

33. Wyss, R.; Grasso, L.; Wolf, C.; Grosse, W.; Demurtas, D.; Vogel, H., Molecular and Dimensional Profiling of Highly Purified Extracellular Vesicles by Fluorescence Fluctuation Spectroscopy. Analytical Chemistry 2014, 86 (15), 7229–7233.

34. Chen, C.; Zong, S.; Wang, Z.; Lu, J.; Zhu, D.; Zhang, Y.; Zhang, R.; Cui, Y., Visualization and intracellular dynamic tracking of exosomes and exosomal miRNAs using single molecule localization microscopy. Nanoscale 2018, 10 (11), 5154–5162.

35. van den Bogaart, G.; Krasnikov, V.; Poolman, B., Dual-Color Fluorescence-Burst Analysis to Probe Protein Efflux through the Mechanosensitive Channel MscL. Biophysical Journal 2007, 92 (4), 1233–1240.

36. Widengren, J.; Rigler, R., Review - Fluorescence correlation spectroscopy as a tool to investigate chemical reactions in solutions and on cell surfaces. Cell Mol Biol 1998, 44 (5), 857–879.

37. Martinez-Moro, M.; Di Silvio, D.; Moya, S. E., Fluorescence correlation spectroscopy as a tool for the study of the intracellular dynamics and biological fate of protein corona. Biophysical Chemistry 2019, 253, 106218.

38. Schwille, P.; MeyerAlmes, F. J.; Rigler, R., Dual-color fluorescence cross-correlation spectroscopy for multicomponent diffusional analysis in solution. Biophys J 1997, 72 (4), 1878–1886.

39. Bacia, K.; Schwille, P., Practical guidelines for dual-color fluorescence cross-correlation spectroscopy. Nat Protoc 2007, 2 (11), 2842–2856.

40. Aalberts, M.; Stout, T. A. E.; Stoorvogel, W., Prostasomes: extracellular vesicles from the prostate. REPRODUCTION 2014, 147 (1), R1–R14.

41. Ronquist, K. G.; Ek, B.; Morrell, J.; Stavreus-Evers, A.; Ström Holst, B.; Humblot, P.; Ronquist, G.; Larsson, A., Prostasomes from four different species are able to produce extracellular adenosine triphosphate (ATP). Biochimica et Biophysica Acta (BBA) - General Subjects 2013, 1830 (10), 4604–4610.

42. Müller, B. K.; Zaychikov, E.; Bräuchle, C.; Lamb, D. C., Pulsed Interleaved Excitation. Biophysical Journal 2005, 89 (5), 3508–3522.

43. Jazani, S.; Sgouralis, I.; Shafraz, O. M.; Levitus, M.; Sivasankar, S.; Pressé, S., An alternative framework for fluorescence correlation spectroscopy. Nature Communications 2019, 10 (1), 3662.

44. Fu, X.; Song, Y.; Masud, A.; Nuti, K.; DeRouchey, J. E.; Richards, C. I., High-throughput fluorescence correlation spectroscopy enables analysis of surface components of cell-derived vesicles. Analytical and Bioanalytical Chemistry 2020, 412 (11), 2589–2597.

45. Patel, G. K.; Khan, M. A.; Zubair, H.; Srivastava, S. K.; Khushman, M. d.; Singh, S.; Singh, A. P., Comparative analysis of exosome isolation methods using culture supernatant for optimum yield, purity and downstream applications. Scientific Reports 2019, 9 (1), 5335.

46. Zheng, Y.; Hasan, A.; Nejadi Babadaei, M. M.; Behzadi, E.; Nouri, M.; Sharifi, M.; Falahati, M., Exosomes: Multiple-targeted multifunctional biological nanoparticles in the diagnosis, drug delivery, and imaging of cancer cells. Biomedicine & Pharmacotherapy 2020, 129, 110442.

47. Liu, C.; Zhang, W.; Li, Y.; Chang, J.; Tian, F.; Zhao, F.; Ma, Y.; Sun, J., Microfluidic Sonication To Assemble Exosome Membrane-Coated Nanoparticles for Immune Evasion-Mediated Targeting. Nano Letters 2019, 19 (11), 7836–7844.

48. Gambin, Y.; Sierecki, E.; Polinkovsky, M.; Alexandrov, K.; Parton, R., Single molecule analysis reveals self assembly and nanoscale segregation of two distinct cavin subcomplexes on caveolae (602.1). The FASEB Journal 2014, 28 (1_supplement), 602.1.

49. Ingargiola, A.; Lerner, E.; Chung, S.; Weiss, S.; Michalet, X., FRETBursts: An Open Source Toolkit for Analysis of Freely-Diffusing Single-Molecule FRET. PLOS ONE 2016, 11 (8), e0160716.

50. Widengren, J.; Kudryavtsev, V.; Antonik, M.; Berger, S.; Gerken, M.; Seidel, C. A. M., Single-Molecule Detection and Identification of Multiple Species by Multiparameter Fluorescence Detection. Analytical Chemistry 2006, 78 (6), 2039–2050.

51. Ambrose, B.; Baxter, J. M.; Cully, J.; Willmott, M.; Steele, E. M.; Bateman, B. C.; Martin-Fernandez, M. L.; Cadby, A.; Shewring, J.; Aaldering, M.; Craggs, T. D., The smfBox is an open-source platform for single-molecule FRET. Nature Communications 2020, 11 (1), 5641.

52. Hagai, D.; Lerner, E., Systematic Assessment of Burst Impurity in Confocal-Based Single-Molecule Fluorescence Detection Using Brownian Motion Simulations. Molecules 2019, 24 (14).

53. Eggeling, C.; Widengren, J.; Rigler, R.; Seidel, C. A. M., Photobleaching of Fluorescent Dyes under Conditions Used for Single-Molecule Detection: Evidence of Two-Step Photolysis. Analytical Chemistry 1998, 70 (13), 2651–2659.

54. Widengren, J.; Rigler, R., Mechanisms of photobleaching investigated by fluorescence correlation spectroscopy. Bioimaging 1996, 4 (3), 149–157.

55. Mitronova, G. Y.; Belov, V. N.; Bossi, M. L.; Wurm, C. A.; Meyer, L.; Medda, R.; Moneron, G.; Bretschneider, S.; Eggeling, C.; Jakobs, S.; Hell, S. W., New Fluorinated Rhodamines for Optical Microscopy and Nanoscopy. Chemistry – A European Journal 2010, 16 (15), 4477–4488.

56. Zade-Oppen, A. M. M., Posthypertonic Hemolysis in Sodium Chloride Systems. Acta Physiologica Scandinavica 1968, 73 (3), 341–364.

