## Supporting information for "Coincident Fluorescence Burst Analysis of dUTP-Loaded Exosome-Mimetic Nanovesicles"

### Figures

**Figure S1.** Schematic of the detection volume and its lateral and radial dimensions; in this setup, the ratio of the axial to lateral dimension was $z_0/w_0 = 5$ and 5.1 for the red and green wavelengths, respectively. 4

### Tables

#### S1- FCCS measurements

##### S1-1- Theory

In simple FCS, the auto-correlation function $G(\tau)$ of the recorded fluorescence fluctuations $\delta F(t) = F(t) - \langle F(t) \rangle$ around the average fluorescence $\langle F(t) \rangle$ is calculated as a function of the time delay $\tau$. Symphotime64 software (PicoQuant) was used for data acquisition and the correlation curve:

$$G_{exp}(\tau) = \frac{\langle F(t)F(t+\tau)\rangle}{\langle F(t)\rangle^2} = 1 + \frac{\langle \delta F(t)\delta F(t+\tau)\rangle}{\langle F(t)\rangle^2} \quad \text{(S1)}$$

For the FCS analysis, the experimental curves for red and green colors were fitted to single and double diffusion functions of the following form, respectively.[1, 2]

$$G_{fit}(\tau) = 1 + \frac{1}{N} \times \sum_i \frac{N_i}{\left(1+\frac{\tau}{\tau_{Di}}\right)\sqrt{\left(1+\left(\frac{w_0}{z_0}\right)^2 \times \frac{\tau}{\tau_{Di}}\right)}} \quad \text{(S2)}$$

where $N = \frac{1}{(\sum_i N_i)^2}$ in which $N_i$ stands for the average number of $i$-th species of fluorescent particles in the confocal volume[2], and the time constant $\tau_{Di}$ is the corresponding average diffusion time through the detection volume. $w_0$ and $z_0$ are the lateral and axial $1/e^2$-radius of the focal detection volume, respectively,[2] as shown in **Figure S1**. The values of $w_0$ for the green and red signal were assessed with separate calibration measurements (see also section **S1-3**).

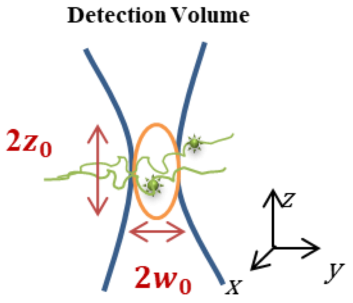

**Figure S1.** Schematic of the detection volume and its lateral and radial dimensions; in this setup, the ratio of the axial to lateral dimension was $\frac{z_0}{w_0}$ =5 and 5.1 for the red and green wavelengths, respectively.

The above expression of the correlation function assumes isotropic particle diffusion and random motion follows the Fick's second law.[3]

Similarly to FCS, in FCCS the cross correlation $G_{cross}(\tau)$ between the fluorescence signals of two species emitting at different wavelengths is calculated from the experimental dual color time traces. Similarly, $G_{cross}(\tau)$ was given as follows: [4]

$$G_{cross}(\tau) = \frac{\langle F_r(t)F_g(t+\tau)\rangle}{\langle F_r(t)F_g(t)\rangle} = 1 + \frac{\langle \delta F_r(t)\delta F_g(t+\tau)\rangle}{\langle F_r(t)F_g(t)\rangle} \quad \text{(S3)}$$

where $F_r(t)$ and $F_g(t)$ stand for the real time traces of the red and green fluorescent signals, respectively. Having determined the average number of red and green particles, $N_r$ and $N_g$, respectively, from the FCS analyses, the average number of two-colored particles ($N_{rg}$) and their average diffusion time ($\tau_{Drg}$) in the confocal volume are determined by fitting the cross correlation curve $G_{cross}(\tau)$ with the following function while ${w_{0(rg)}}^2 = \frac{{w_{0(red)}}^2 + {w_{0(green)}}^2}{2}$. [5]

$$G_{rgFIT}(\tau) = 1 + \frac{N_{rg}}{N_r N_g} \times \frac{1}{\left(1+\frac{\tau}{\tau_{Drg}}\right)\sqrt{\left(1+\left(\frac{w_{0(rg)}}{z_0}\right)^2 \times \frac{\tau}{\tau_{Drg}}\right)}} \quad \text{(S4)}$$

#### S1-2- Fluorescent dyes

a) **Green dye for samples:** Chroma Tide™ Alexa Fluor™ 488-5-dUTP, Thermo-fisher, No. C11397
b) **Red dye for samples:** CellVue® Claret Far Red Fluorescent Cell Linker Kit, Sigma-Aldrich, No. cb_000730
c) **Green dye for Calibration and detection volume measurement:** N-Fmoc Rhodamine 110, Sigma-Aldrich, No. 74171
d) **Red dye for Calibration and detection volume measurement:** Cy5 dye, Thermo-fisher.

#### S1- 3-Calibration and confocal volume measurement

The single color auto correlation curves of three control dyes namely Rhodamine110 and Cy5 at concentrations of approximately 2 nM, and at non-saturating excitation intensities (13.4$\mu$W and 20$\mu$W, respectively) were evaluated and their corresponding parameters such as diffusion time ($\tau_D$) or average count rates have been determined and used as reference. By fitting experimental correlation curves obtained for Rhodamine110 and Cy5 in aqueous solution, considering the reported diffusion coefficient values (D) in Ref. 3, the detection volume and $\frac{z_0}{w_0}$ values were estimated for detection in the green and the red emission range, respectively, and at temperature of the lab (T=23.5 $^0$C). According to the estimated confocal volumes and $\frac{z_0}{w_0}$, the lateral $1/e^2$-radii of the detection volume for green and red signals were estimated to be $w_{0(green)} = 254\ nm$ and $w_{0(red)} = 319\ nm$, respectively.[4] The summary of corresponding values is given at **Table S1**.

**Table S1.** Summary of calibration parameters for FCCS setup, measured for Rhodamine110 and Cy5 dyes.

| FCCS Calibration parameters | | | | |
|---|---|---|---|---|
| Dye | D $(cm^2.s^{-1})$ at T=23.5$^0$C | Confocal Volume (fL) | $\frac{z_0}{w_0}$ | $w_0\ (nm)$ |
| Rhodamine110 | $4.515 \times 10^{-6}$ | 0.165 | 5.1 | 254 |
| Cy5 | $3.458 \times 10^{-6}$ | 0.320 | 5.0 | 319 |

The average count rate of dUTP.Alexa488 dye was measured at the same excitation power (80$\mu$W) and 20MHz repetition rate as the FCCS experiments on the main samples and during similar measurement time (>1hour) at 1 nM concentration. By fitting the corresponding correlation curve to a single diffusion function (see eq. **S1**), the average count rate of single dUTP.Alexa488 molecule was estimated as ≃9.5 kcounts/s.

#### S1- 4-FCCS analysis

For first estimation the experimental auto and cross correlation curves were fitted to the diffusion functions of section **S1-1**. The average diffusion times of CellVue, dUTP.Alexa488 and two colored vesicles ($\tau_r, \tau_g$ and $\tau_{rg}$) and their average numbers ($N_r, N_g, N_{rg}$) in detection volume were inferred from a standard FCCS analysis. The measured G-curves for the red auto- and the red-green cross-correlation curves were fitted to a single diffusion function while to obtain good agreement the green correlation curves had to be fitted to the sum of two diffusion functions. The fits are shown in **Figures S2a-f**. The **Table S2** provides the values extrapolated from the fits. As shown in this table, the second diffusion time for green G-curve fit ($\tau_{g2}$) is much smaller than the first one, which is similar to the diffusion times of free dye molecules (Alexa488) just showing slight longer value since it is attached to dUTP cargo.

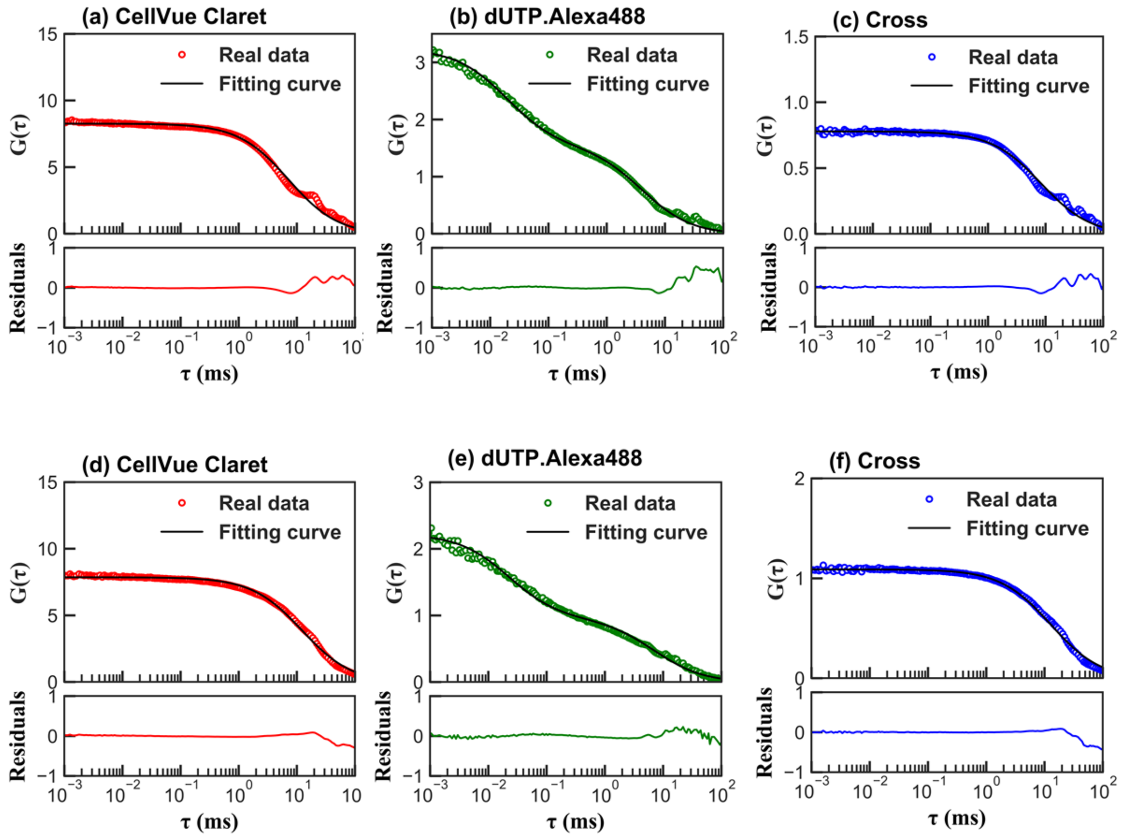

**Figure S2.** (a)-(c) and (d)-(f) The correlation curves of CellVue Claret, dUTP.Alexa488 and their dual colors and their fitting functions, respectively measured for loaded Exo and RBC nanovesicles. Plots of their corresponding fitting residuals also are show below each curve.

**Table S2.** FCCS analysis results: Estimated average values of loading yields ($\boldsymbol{\eta_{ave}}$), diffusion times of nanovesicles ($\boldsymbol{\tau_r}$), dUTP molecules ($\boldsymbol{\tau_g}$) and dUTP-loaded nanovesicles ($\boldsymbol{\tau_{rg}}$) and corresponding average sizes for (a) EXO and (b) RBC samples, respectively. (c) The average diffusion times extracted from fitting results of correlation curves.

| | (a) EXO | | | | |
|---|---|---|---|---|---|
| G curve | $\boldsymbol{\tau_D}$ ms | $R$ nm | $\boldsymbol{N}$ | $\boldsymbol{\tau_{D2}}$ ms | $\boldsymbol{N_2}$ |
| Red | $\tau_r = 7.26$ | $R_r = 62$ | $N_r = 0.12$ | - | - |
| Green | $\tau_g = 4.06$ | $R_g = 54$ | $N_g = 0.15$ | $\tau_{g2}$ =0.02 | 0.16 |
| Cross | $\tau_{rg} = 8.44$ | $R_{rg} = 88$ | $\frac{N_{rg}}{N_r N_g} = 0.78$ | - | - |
| Loading Yield | $\eta_{ave} = \frac{N_{rg}}{N_r N_g} \times N_g = 0.78 \times 0.15 = 11.7\%$ | | | | |

| | (b) RBC |
|---|---|

| G curve | $\boldsymbol{\tau_D}$ ms | $R$ nm | $\boldsymbol{N}$ | $\boldsymbol{\tau_{D2}}$ ms | $\boldsymbol{N_2}$ |
|---|---|---|---|---|---|
| Red | $\tau_r = 12.45$ | $R_r = 106$ | $N_r = 0.13$ | - | - |
| Green | $\tau_g = 7.03$ | $R_g = 94$ | $N_g$ =0.20 | 0.02 | 0.25 |
| Cross | $\tau_{rg} = 12.97$ | $R_{rg} = 135$ | $\frac{N_{rg}}{N_r N_g} = 1.09$ | - | - |
| Loading Yield | $\eta_{ave} = \frac{N_{rg}}{N_r N_g} \times N_g = 1.09 \times 0.20 = 21.8\%$ | | | | |

According to the average number of particles obtained from the green auto- ($N_g$) and the cross-correlation curves, the average dUTP-loading yield can be calculated as $\eta_{ave} = \frac{N_{rg}}{N_r N_g} \times N_g$. Based on FCCS G-curves, the average loading yield for RBC sample was obtained beyond 100%, which is impossible and is due to aggregation and existence of free green molecules in solution. Moreover, according to the average diffusion times, the average radius size of red vesicles ($R_r$), green particles ($R_g$), and two-colored vesicles ($R_{rg}$) have been estimated and summarized in **Table S2**, showing relatively large values beyond the reasonable size range of exosomes which is less than 75nm in radius.

#### S2- AFM Measurement

To confirm the vesicle shape and size of the samples, their morphology and size distribution were systematically checked using Atomic Force Microscopy (AFM).

##### S2-1- AFM images

Examples of typical images obtained by AFM on EXO and RBC nanovesicles are shown in **Figures S3** and **S4**. In tapping mode, the cantilever is oscillating close to the surface. The AFM amplitude images are obtained from the amplitude of such oscillations. The phase images map the phase shift of such mechanical oscillations with respect to the AFM driving signal[6]. The phase shift can reflect various properties of the surface, such as their viscoelasticity or softness. The phase images (**S3c** and **S4c**) provide clearer pictures of vesicles. They also indicate that agglomeration is more pronounced in RBC samples.

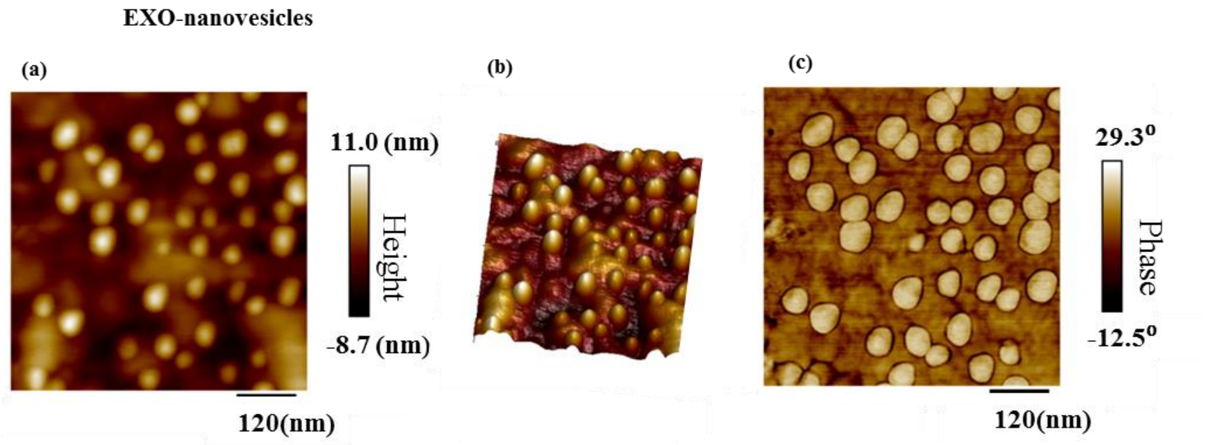

**Figure S3.** AFM images of EXO nanovesicles on area of (1$\mu$m)$^2$: (a) height image, (b) same as (a) in 3D, and (c) phase image.

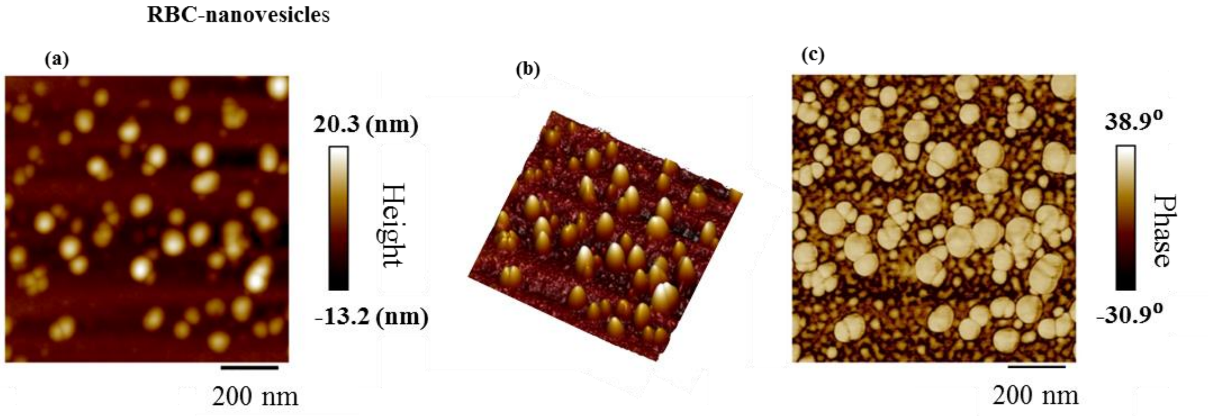

**Figure S4.** AFM images of RBC nanovesicles on area of (1$\mu$m)$^2$: (a) height image, (b) same as (a) in 3D, and (c) phase image.

#### S2-2- AFM Data analysis

The size distributions of all the vesicles were quantified statistically by taking larger AFM images with a typical scan size of (5 μm)$^2$ for each sample. The two-dimensional AFM images were analyzed using the 'imfindcircle' function in the image processing package of a commercial software (Mathworks ®). The software detected the location of the nanovesicles and fitted each of them to a circular shape. The fit was used to extrapolate the individual vesicle radius (R) and then to plot size distribution histograms of the nanovesicles (**Figure.2** main article). The image analysis of the AFM data obtained on EXO and RBC nanovesicles over an area of $(5\mu m)^2$ is illustrated by **Figure S5**.

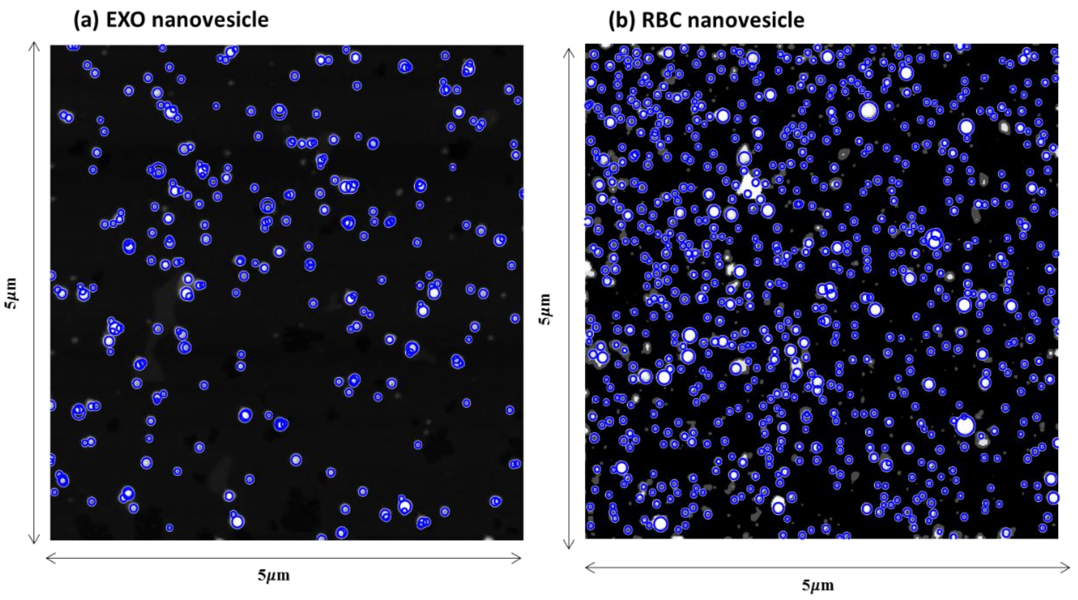

**Figure S5.** Image processing of AFM height image of (a) EXO and (b) RBC nanovesicles on an area of $(5\mu m)^2$.

For comparison, the size histograms of semEV and EXO vesicles were also measured. They are shown in **Figure S6,** which clearly shows a shift to larger sizes and broader distribution of the EXO nanovesicles, when compared to semEV.

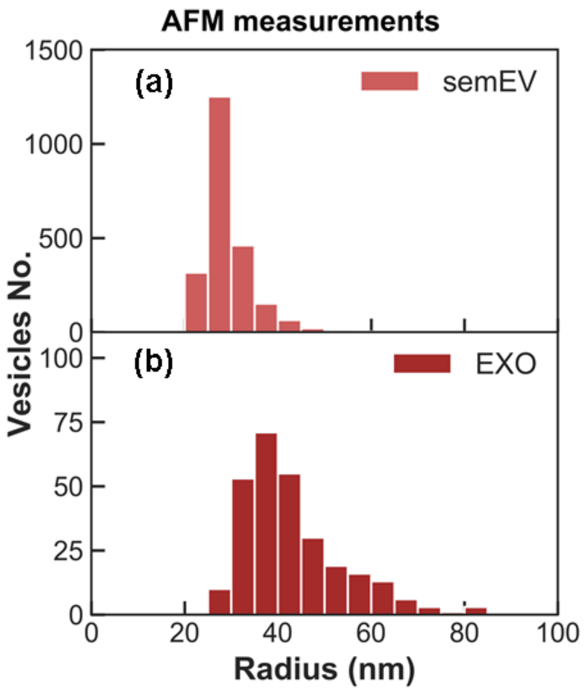

**Figure S6.** AFM size histograms of (a) semEV and (b) EXO. The EXO vesicles show a shift to larger sizes and have a broader distribution compared to semEV.

### S3- DC-CFB Data analysis

#### S3-1- Time-gating and lifetime histograms

The absorption and emission spectra of Alexa488 and CellVue Claret dyes and corresponding spectral ranges of applied detection filters are shown with shaded windows in **Figure S7**, depicting no spectral overlap between red and green channels.

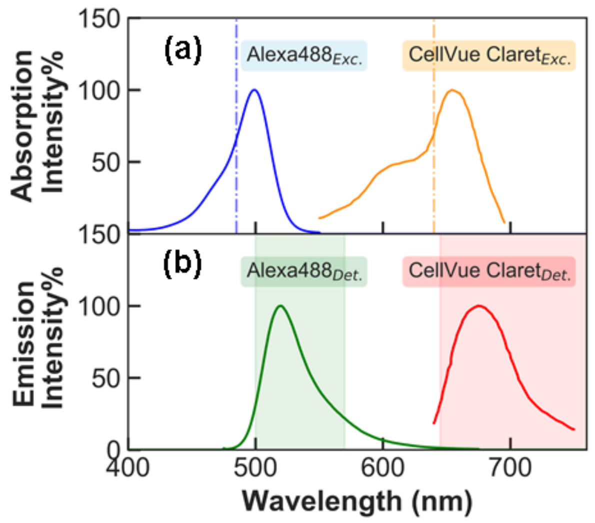

**Figure S7.** The absorption (a) and emission (b) spectra of Alexa488 (data from Thermofisher[7]) and CellVue Claret (data from Sigmaaldrich[8]) dyes. The blue and orange vertical lines in (a) show the excitation laser wavelengths while the green and red shaded windows in (b) depict the detected spectral ranges of corresponding channels.

With the help of time correlated single photon counting (TCSPC)[4] modules and using the Symphotime 64 software (Picoquant), the photon streams were collected by a minimal binning time equal to 16 ps. For coincident fluorescence burst analysis, the photon timestamps in the red and green channel were first time-gated. To find an appropriate time gating window, the lifetime histograms of the photon counts of the green and red detectors were plotted, as shown in **Figure S8**. The green and red timestamps were selected during green- and red-shaded windows, corresponding to the pulsed interleaved excitation (PIE) sequences of the two excitation lasers and the fluorescence lifetimes of the two dyes[9]. Thereby, cross-talk was suppressed, in particular of green fluorescence into the red channel.

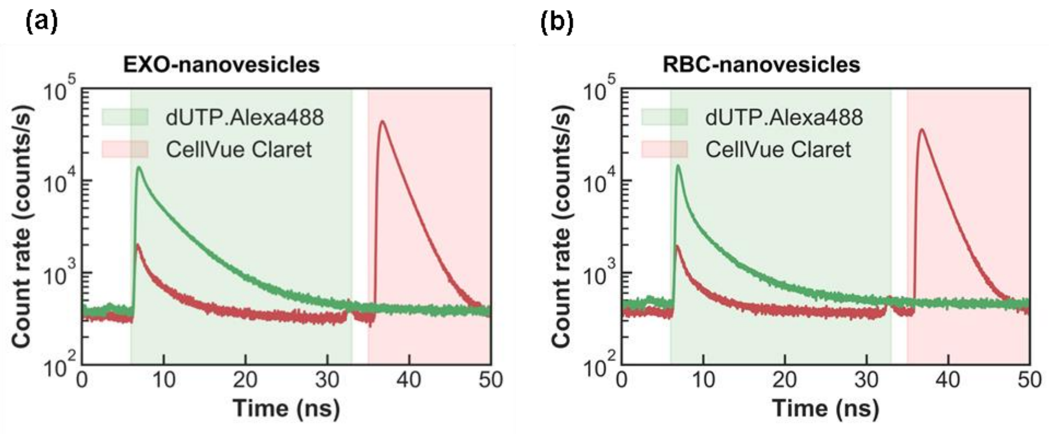

**Figure S8.** Lifetime histograms of CellVue Claret and dUTP.Alexa488 and their corresponding time-gating windows shaded in red and green, respectively, for loaded (a) EXO and (b) RBC nanovesicles.

#### S3-2- Background Rates

Before searching for the fluorescence bursts, it is essential to evaluate the background rate (BG-rate) in order to estimate an appropriate threshold for the burst search, and to correct the raw burst counts by subtracting time-varying background counts. These might occur for several reasons, such as detector dark counts, after pulsing, out-of-focus molecules, photo-bleaching or evaporation of the sample solution[5]. The time-dependent BG-rates in the red and green channels were estimated in time windows of 50s, by fitting the distribution of inter-photon delays using the Maximum Likelihood Estimation (MLE) method, embedded in the open source code FRETBurst[9]. Typical BG-rate for a measurement longer than 50 minutes duration for the red and green channels are shown in **Figure S9**. Both sets of measurements indicate an increase of about 0.5 (kCounts/s) in BG-rates within 50 minutes imposing the need to perform this calibration before any CFB data analysis.

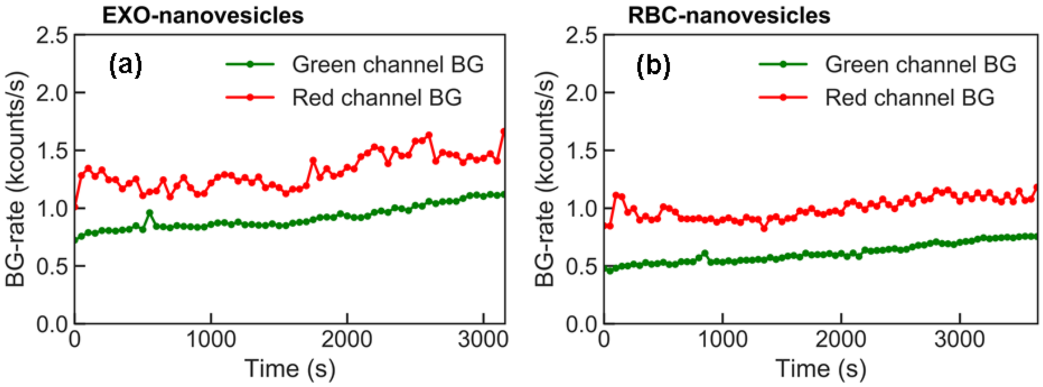

**Figure S9.** BG-rates of red and green channels versus time in 50s time windows, during FCCS measurement time for loaded (a) EXO nanovesicles and (b) RBC nanovesicles. There is slight increase in the background count rate in time.

#### S3-3- Burst Conditions

All the bursts in the red and green photon streams were identified using a "sliding window" algorithm.[9] In this method, the code searches for $m$ consecutive photons detected at a count rate larger than $F * BG.rate$ and during a period shorter than $\Delta t = (m-1)/(F * BG.rate)$.[9] Two parameters $m$ and $F$ are of critical importance for burst search. Due to the lack of a universal criterion to choose these values, appropriate validation routines had to be established. Specifically to identify optimal $m$ and $F$ values (1) for red burst analysis, we performed systematic comparisons with AFM measurements and (2) for green burst analysis, we compared the *Coincident*-red and *Coincident*-green histograms as explained in the text.

##### a) AFM-RFB Comparison

By comparing the normalized size histograms retrieved from AFM measurements and RFB in **Figure S10**. Both AFM and red burst method show similar shape around the peak but their standard deviations are different which is attributed to the agglomeration of vesicles at larger sizes, since we have diluted the sample solutions much more for AFM measurements (around $10^{-11}$ dilution).

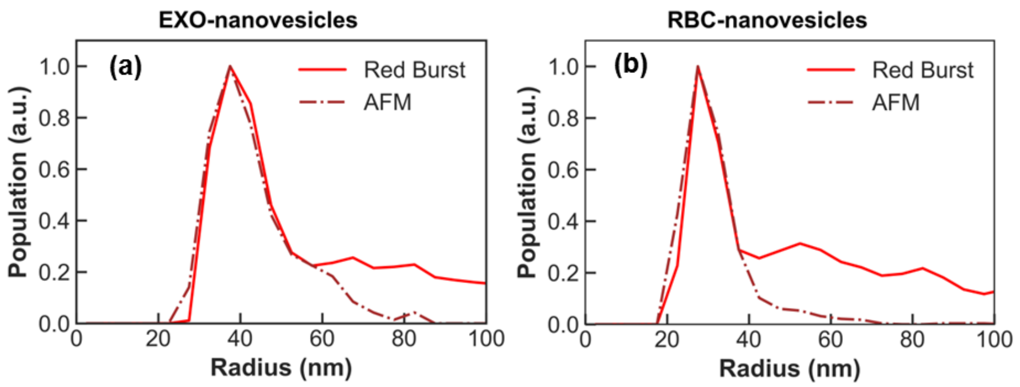

**Figure S10.** Comparing the normalized size distributions obtained from AFM and red bursts for (a) EXO nanovesicles and (b) RBC nanovesicles. The red sloid curves depict red bursts distributions and brown dash-dotted curves represent AFM distributions.

In **Table S3**, is a comparative study between the size histograms retrieved from AFM and RFB for EXO and RBC nanovesicles. Additionally, as Co-red Bursts columns we show the results of the *coincident*-red burst analysis (see also next section) corresponding to loaded vesicles. Since the distribution shapes of AFM and red bursts are right-skewed, the values of mode, median and interquartile range (IQR) of each histogram are reported in this table.

**Table S3.** The summary of the comparison between the size histograms obtained from AFM, RFB, and coincident red bursts corresponding to loaded vesicles (see also **S3-3b**) for EXO and RBC nanovesicles.

| Sample | EXO | | | RBC | | |
|---|---|---|---|---|---|---|
| **Parameters (nm)** | **AFM** | **Red Bursts** | **Co-red Bursts** | **AFM** | **Red Bursts** | **Co-red Bursts** |
| $R_{max}$ | 37 | 37 | 60 | 27 | 27 | 42 |
| $\bar{R} \pm std$ | 34±3 | 46±12 | 55±6 | 29±4 | 42±15 | 49±9 |
| $mode$ | 27 | 32 | 49 | 25 | 26 | 38 |
| $Median \pm IQR$ | 34±4 | 46±34 | 56±9 | 29±6 | 42±15 | 47±11 |

#### b) Histogram checks between coincident red and coincident green bursts

Identifying coincident fluorescence bursts in the stream of red and green photons is one of the most crucial steps in these analyses. In this step, the start and stop times of each red burst are compared to the ones in all green bursts, to find the bursts which have an overlap in time. The analysis allows for a central delay between overlapping red and green bursts ($\Delta t_{rg}$) of at most 5µs, as shown in **Figure S11**.

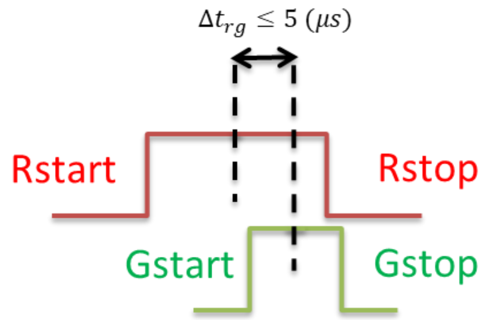

**Figure S11.** The coincidence check between overlapping red and green bursts, such as their central delay could be less than $5\mu$s.

The maximum value of the central time delay between overlapping red and green bursts is chosen by comparing the normalized corresponding size histograms of coincident red bursts and coincident green bursts defined as: $H_m^{norm}(R) = \frac{H_m(R)}{max\{H_m(R)\}}$, where $H_m(R)$ is the distribution of the *coincident*-red or *coincident*-green bursts, respectively, with $m = Co - red$ or $Co - green$. The optimal values of $F_g$, $m_g$ and $\Delta t_{rg}$ are determined by minimizing the difference between *coincident*-red and *coincident*-green normalized histograms that is: $\varepsilon(R) = H_{Co-red}^{norm}(R) - H_{Co-green}^{norm}(R)$ in the range of $R \leq 75\ nm$, as shown in **Figure S12**. This difference is determined to be less than 0.26 and 0.37 with average of about 0.03 and 0.08 for EXO and RBC samples, respectively.

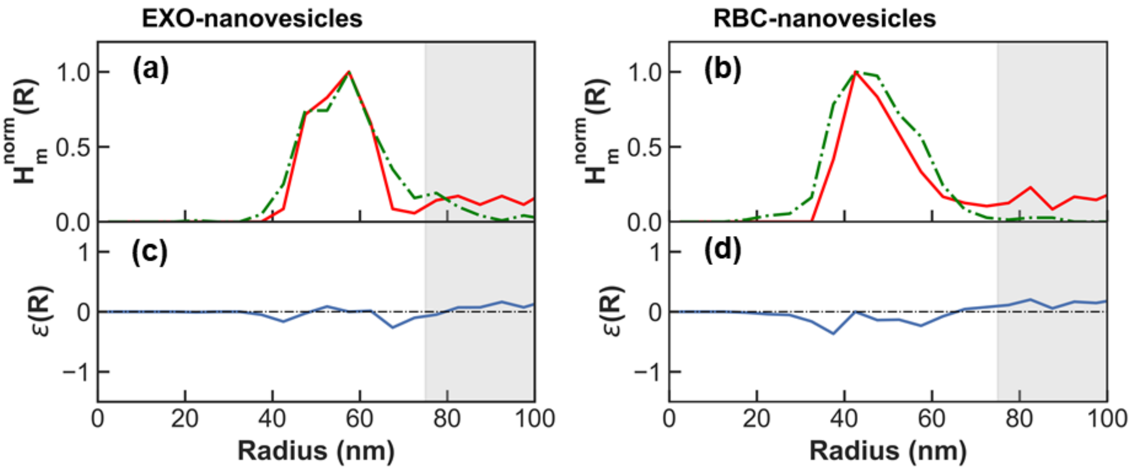

**Figure S12.** Plots of normalized histograms of coincident red (solid red) and coincident green (dash-dotted green) bursts for (a) EXO and (b) RBC nanovesicles, (c) and (d) their difference.

The parameters yielding the best agreement are given in **Table S4**, together with corresponding maximum and average error.

**Table S4.** The optimized parameters for red and green burst searches for EXO and RBC nanovesicles and corresponding maximum and average errors in the range of $R \leq 75\ nm$.

| Sample | EXO-nanovesicles | | RBC-nanovesicles | |
|---|---|---|---|---|
| Parameters | Red | Green | Red | Green |
| $\boldsymbol{F}$ | 6 | 5 | 7 | 5 |
| $\boldsymbol{M}$ | 36 | 19 | 23 | 11 |
| $\boldsymbol{max}\{\boldsymbol{\varepsilon}(\boldsymbol{R})\}$ | 0.26 | | 0.37 | |
| $\langle\boldsymbol{\varepsilon}(\boldsymbol{R})\rangle$ | 0.03 | | 0.08 | |

The size histograms of red bursts, green bursts, and the coincident red bursts corresponding to loaded nanovesicles, are plotted in **Figure S13**, for EXO and RBC nanovesicles.

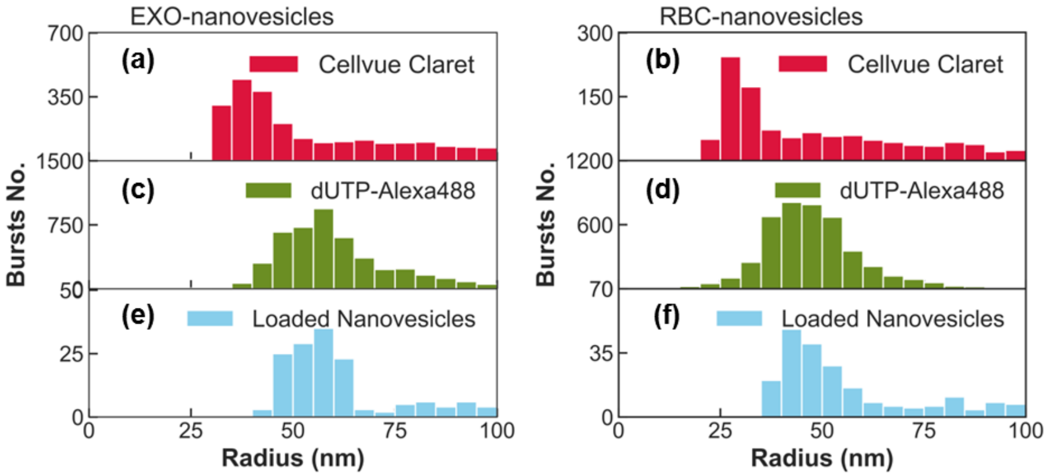

**Figure S13.** Size histograms of red bursts (a, d), green bursts (b, e), and the coincident red bursts (c, f) corresponding to the loaded nanovesicles, for EXO and RBC nanovesicles, respectively.

To estimate the number of cargos per vesicle, the count rate of coincident green bursts were divided by the average count rate of single dUTP-Alexa488 cargo molecule. **Figure S14** depicts the histograms of the number of cargo molecules per vesicle, showing most population in the range of less than 2.75. The average number of cargo molecules per vesicle in this range was 1.50 and 1.15 for EXO and RBC samples, respectively.

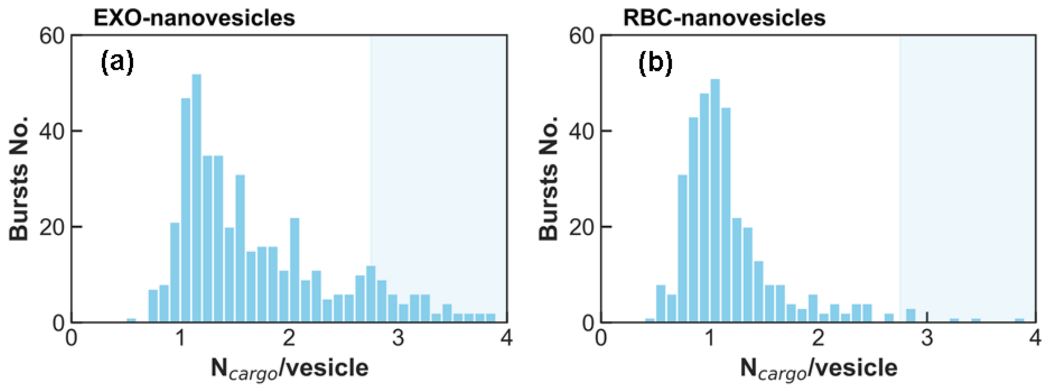

**Figure S14.** Histograms of number of cargo molecules per vesicle in loaded ones for (a) EXO and (b) RBC samples.

#### S3-4- Photobleaching effects

According to reported values for the absorption quantum yield ($\phi_{abs}$) and photobleaching cross section ($\sigma_{bl}$) of the Alexa488 dye at high excitation powers,[10] thephotobleaching rate of this dye can be estimated as $k_z = 17.72\ s^{-1}$ by one order of magnitude less in the low excitation power regimen used for all experiments. The latter corresponds to ~40 kW/cm$^2$ excitation intensities for the green channel, at which the dye is below the saturation regime and the upper singlet and triplet states would have very low probability to be populated with an ensuing negligible role in photobleaching. More specifically, the corresponding photobleaching quantum yield is in the order of $10^{-6}$. The percentage of nanovesicles potentially affected by photobleaching within their diffusion time in the confocal excitation volume can be estimated as $c = e^{-k_z \cdot \tau_{Co-Gburst}}$ where $\tau_{Co-Gburst}$ denotes the temporal duration of the coincident-green bursts.[11] The histogram of the population percentage of unbleached nanovesicles for loaded EXO and RBC nanovesicles inferred by further analysis of the experimental data is plotted in **Figure S15**.

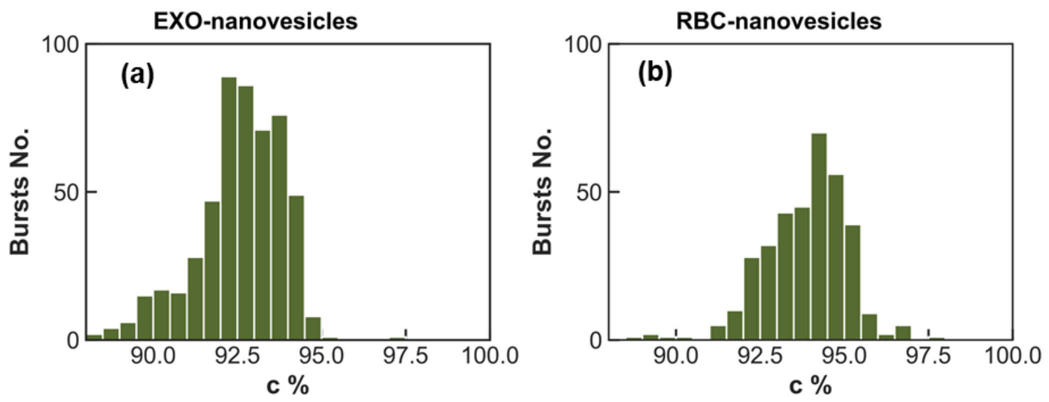

**Figure S15.** Histograms of the percentage of intact remained vesicles to photobleaching within passage time for EXO (a) and RBC (b) loaded nanovesicles.

#### References

1. Widengren, J.; Mets, Ü., Conceptual Basis of Fluorescence Correlation Spectroscopy and Related Techniques as Tools in Bioscience. In *Single Molecule Detection in Solution*, 2002; pp 69-120.
2. Fluorescence Correlation Spectroscopy. In *Principles of Fluorescence Spectroscopy*, Lakowicz, J. R., Ed. Springer US: Boston, MA, 2006; pp 797-840.
3. Banks, D. S.; Fradin, C., Anomalous Diffusion of Proteins Due to Molecular Crowding. *Biophysical Journal* **2005,** *89* (5), 2960-2971.
4. https://www.picoquant.com/products/category/software/symphotime-64-fluorescence-lifetime-imaging-and-correlation-software.
5. Bacia, K.; Kim, S. A.; Schwille, P., Fluorescence cross-correlation spectroscopy in living cells. *Nature Methods* **2006,** *3* (2), 83-89.
6. Forchheimer, D.; Forchheimer, R.; Haviland, D. B., Improving image contrast and material discrimination with nonlinear response in bimodal atomic force microscopy. *Nature Communications* **2015,** *6* (1), 6270.
7. https://www.thermofisher.com/order/fluorescence-spectraviewer#!/ https://fretbursts.readthedocs.io/en/latest/.
8. https://www.sigmaaldrich.com/SE/en/product/sigma/minclaret?gclid=CjwKCAjw0qOIBhBhEiwAyvVcfxQ3nVA7CQtxDc1d2BMq9gzTvLsI9wkkt7nY0pDgtMfzlcXBBRxfvBoCF4sQAvD_BwE.
9. Ingargiola, A.; Lerner, E.; Chung, S.; Weiss, S.; Michalet, X., FRETBursts: An Open Source Toolkit for Analysis of Freely-Diffusing Single-Molecule FRET. *PLoS One* **2016,** *11* (8), e0160716-e0160716.
10. Mitronova, G. Y.; Belov, V. N.; Bossi, M. L.; Wurm, C. A.; Meyer, L.; Medda, R.; Moneron, G.; Bretschneider, S.; Eggeling, C.; Jakobs, S.; Hell, S. W., New Fluorinated Rhodamines for Optical Microscopy and Nanoscopy. *Chemistry – A European Journal* **2010,** *16* (15), 4477-4488.
11. Eggeling, C.; Widengren, J.; Rigler, R.; Seidel, C. A. M., Photobleaching of Fluorescent Dyes under Conditions Used for Single-Molecule Detection:  Evidence of Two-Step Photolysis. *Analytical Chemistry* **1998,** *70* (13), 2651-2659.
